# Single-molecule imaging of cytoplasmic dynein *in cellulo* reveals the mechanism of motor activation and cargo movement

**DOI:** 10.1101/2021.04.05.438428

**Authors:** Nireekshit Addanki Tirumala, Gregory Redpath, Sarah Viktoria Skerhut, Pritha Dolai, Natasha Kapoor-Kaushik, Nicholas Ariotti, K Vijay Kumar, Vaishnavi Ananthanarayanan

## Abstract

Cytoplasmic dynein 1 (dynein) is the primary minus end-directed motor protein in most eukaryotic cells. Dynein remains in an inactive conformation until the formation of a tripartite complex comprising dynein, its regulator dynactin and a cargo adaptor. How this process of dynein activation occurs is unclear, since it entails the formation of a three-protein complex inside the crowded environs of a cell. Here, we employed live-cell, single-molecule imaging to visualise and track fluorescently tagged dynein. First, we observed that only ~30% of dynein molecules that bound to the microtubule (MT) engaged in minus end-directed movement, and that too for a short duration of ~0.6 s. Next, using high-resolution imaging in live and fixed cells, and using correlative light and electron microscopy, we discovered that dynactin and endosomal cargo remained in proximity to each other and to MTs. We then employed two-colour imaging to visualise cargo movement effected by single motor binding. Finally, we performed long-term imaging to show short movements are sufficient to drive cargo to the perinuclear region of the cell. We then used these discoveries as the basis for a stochastic model incorporating dynamic motors binding to cargo located along MTs, and also developed a coarse-grained 3-state run-and-tumble particle (RTP) model for the cargo that quantitatively recapitulates the emergent statistics of cargo movement. Taken together, we discovered a search mechanism that is facilitated by dynein’s frequent MT binding-unbinding kinetics: (1) in a futile event when dynein does not encounter cargo anchored in proximity to the MT, dynein dissociates and diffuses into the cytoplasm, (2) when dynein encounters cargo and dynactin upon MT-binding, it moves cargo in a short run. Several of these short runs are undertaken in succession for long-range directed movement. In conclusion, we demonstrate that dynein activation and cargo capture are coupled in a step that relies on the reduction of dimensionality to enable minus end-directed transport *in cellulo*, and that complex cargo behaviour emerges from stochastic motor-cargo interactions.

## Introduction

The dynein family of motor proteins comprises axonemal and cytoplasmic dyneins. While cytoplasmic dynein 2 plays a critical role in intraflagellar transport (Höök & Vallee, 2006), cytoplasmic dynein 1 (dynein henceforth) is responsible for force production and minus-end directed movement of a variety of cargo in cells containing MTs (Allan, 2011). Dynein is a large complex of homodimers comprising 500kDa heavy chains, and other accessory proteins including light chains, light intermediate chains and intermediate chains, which mediate dimerisation of dynein and thereby its processivity. The activity of motor proteins is typically regulated: several kinesins assume an autoinhibited conformation until attachment to cargo (Verhey & Hammond, 2009); dynein was first found to be regulated for its processivity by the multi-subunit complex, dynactin (Verhey & Hammond, 2009). More recent studies have additionally implicated cargo adaptors -which link dynein to a multitude of cargo - in the activation of dynein (Vallee *et al*, 2012; Cianfrocco *et al*, 2015; Ananthanarayanan, 2016; Canty & Yildiz, 2020). Dynactin is a large multi subunit complex (Schroer, 2004) that was first identified as an activator of minus-end directed motility of vesicles (Gill *et al*, 1991). Further research indicated that an intact dynactin complex was necessary for dynein’s function and that dynactin could interact with MTs via its p150 subunit (Quintyne *et al*, 1999; Valetti *et al*, 1999). The N-terminal CAP-Gly domain on p150 was also found to be able to interact with growing MT plus ends via EB1/CLIP-170 pathway (Vaughan *et al*, 1999; Watson & Stephens, 2006) and influence intracellular transport (Vaughan *et al*, 2002). However, the interaction between dynein and dynactin, as probed from coimmunoprecipitation assays, was observed to be weak, and over expressing the N-terminal (cytoplasmic) fragment of the cargo adaptor BiCD2 was found to significantly increase dynein-dynactin interaction (Splinter *et al*, 2012).

Cargo adaptors are proteins that link membranous cargo to the motor (Hoogenraad & Akhmanova, 2016), with specific cargo adaptors being unique to different types of cargo (Reck-Peterson *et al*, 2018; Olenick & Holzbaur, 2019). Recent single molecule *in vitro* research has established that formation of the dynein-dynactin-cargo adaptor (DDC) complex is essential for processive motion (McKenney *et al*, 2014; Schlager *et al*, 2014). Cryo-EM studies later reveaíed that the formation of the DDC complex relieved the autoinhibition of dynein and reoriented the motor for processive movement (Torisawa *et al*, 2014; Zhang *et al*, 2017; Chowdhury *et al*, 2015). The DDC complex ameliorates the force produced by single dynein motors from 1pN to about 6pN (Belyy *et al*, 2016). Thus, our current understanding suggests that formation of DDC complex is essential in dynein driven transport. While it is clear that the tripartite complex formation is an essential first step in the activation of the dynein motor, it is unknown how this process occurs in a living cell, amid its crowded environs and the independent dynamics of each component of the tripartite complex. Here, we employ several strategies including single-molecule imaging, correlative light and electron microscopy (CLEM), high-resolution fluorescence microscopy and stochastic modelling to reveal the mechanism of formation of the tripartite complex and therefore, activation of dynein.

## Results

### Dynein interacts transiently with the MT

To probe the kinetics of dynein in living cells, we employed HeLa cells stably expressing mouse dynein heavy chain (DYNC1H1) tagged with GFP (mDHC-GFP, (Poser *et al*, 2008)). To visualise mDHC-GFP, we adapted highly inclined and laminated optical sheet (HILO) microscopy (Tokunaga *et al*, 2008)), Fig. S1a). When cells expressing low levels of mDHC-GFP were observed under a spinning disk confocal (SD) microscope, the fluorescence signal appeared cytosolic, with no discernible dynein punctae. However, when the same cells were observed using our modified HILO microscopy, distinct fluorescent spots were visible (Fig. S1bi, ii).

We adapted our microscopy protocol to obscure dynein diffusing in the cytoplasm and to only observe dynein that was bound to and resided on the MT ((Ananthanarayanan *et al*, 2013; Ananthanarayanan & Tolić, 2015), Fig. 1a, see SI Methods). We observed that dynein spots appeared afresh and remained in the field of imaging for a short duration (Fig. 1b, Movie S1). We intuited that these corresponded to events where single dynein molecules previously diffusing in the cytoplasm bound to MTs (Fig. S1c-e).

**Fig. 1.**
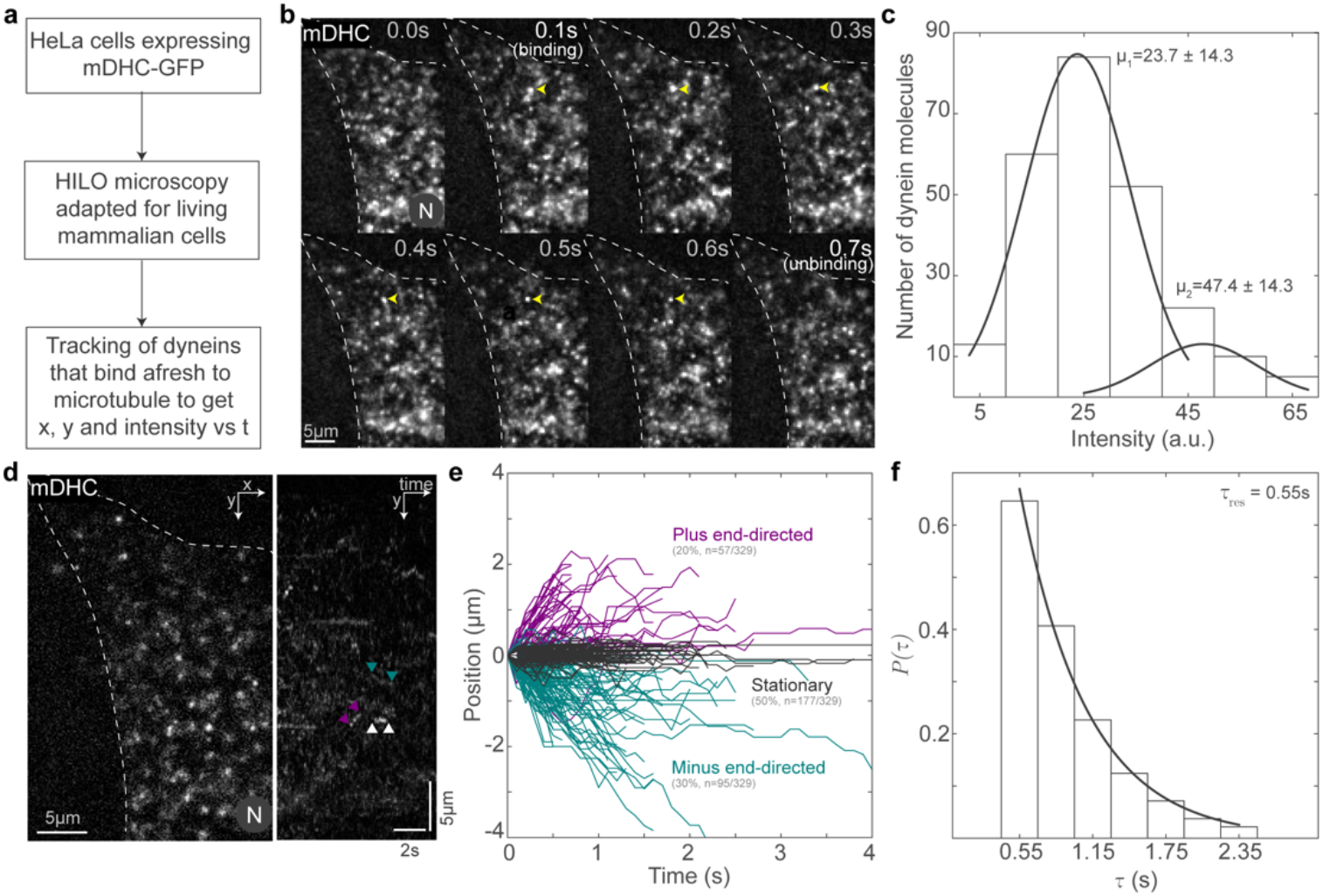
Visualisation of single molecules of dynein in living cells. **a**, Schematic of the protocol followed for the visualisation of single molecules of dynein. **b**, Montage of HILO images showing representative binding and unbinding events of a single fluorescent mDHC molecule (‘(binding)’ and ‘(unbinding)’). The single molecule is indicated with yellow arrowheads for the duration of the time it remains bound in the field of view. Time is indicated at the top right of each image in the montage. **c**, Intensity histogram of single molecules of dynein with the Gaussian fits (grey line). The mean ± s.d. of the Gaussian distributions is indicated above the fits. **d**, HILO image (left) and kymograph (right) of a cell expressing mDHC-GFP. Representative stationary, minus end-directed and plus end-directed events are indicated with the white, teal and magenta arrowheads respectively in the kymograph. **e**, Plot of position vs. time for the single-molecule events tracked, showing stationary events (grey), minus end-directed events (teal) and plus end-directed events (magenta). **f**, Histogram with the residence time of dynein on the MT on the x axis and *P*(*τ*)=1-cumulative frequency on the y axis. The exponential fit (grey line) gave a mean residence time 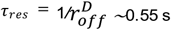. In **b** and **d**, ‘N’ marks the location/direction of the nucleus.

To confirm that the appearance of fluorescent signal on the MT corresponded to binding of a single molecule of dynein to the MT, we analysed the intensity of these fluorescent spots (Fig. 1c). For single dynein molecules, we would expect the intensity histogram to fit to a sum of two Gaussian distributions, one corresponding to a GFP fluorescing from one DHC and the other corresponding to two GFPs fluorescing from both DHCs in the genetic background of these cells. The former primarily arose due to photobleaching of GFP during the course of imaging. Accordingly, the intensity histogram of these fluorescent spots revealed that we indeed observed single dynein molecules since the intensity histogram fit best to a sum of two Gaussians, with the mean of the first Gaussian profile being half that of the second (Fig. 1c).

Next, we analysed the two-dimensional position (*x* and *y*) versus time (*t*) of single molecules of dynein that bound afresh from the cytoplasm to the MT (Fig. 1d). Based on automated thresholding (see SI Methods), we classified the tracks as stationary, minus end-directed and plus end-directed. The plus ends of the MT were predominantly at the periphery in these elongated cells (Fig. S1f), and hence we annotated movement towards the cell center as minus end-directed, and movement away as plus end-directed. We observed that ~50% of all the dynein molecules tracked (n=177/329, N=3 independent experiments from >50 cells) remained stationary upon MT binding, while ~30% (n=95/329) moved towards the minus end (Fig. 1e). The remaining ~20% moved towards the plus end and these arose likely due to attachment of dynein to cargo being moved to the plus end by kinesins (Fig. 1e). The velocity measured for minus-end directed movement of single dyneins was 1.2±0.7 *μ*m/s (mean±s.d.), similar to values reported for mammalian dynein previously (Flores-Rodriguez *et al*, 2011; Zajac *et al*, 2013). We also confirmed that the underlying MT was stable, did not undergo sliding, and therefore did not contribute to the dynein behaviour we observed (Fig. S1g). We then measured the mean residence time (*τ*_res_) of dynein on the MTs to be 0.55 s (95% confidence interval (CI): 0.51-0.59 s) (Fig. 1f). The unbinding rate of dynein from MT 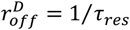 was 1.7 s^-1^, which is similar to the previously reported unbinding rate of single dyneins (Kunwar *et al*, 2011). We verified that this short residence time of dynein on MTs was a true representation of the duration of time that dynein remained attached to the MT and not convolved by GFP’s photobleaching time (Fig. S1h, i). Further, by knocking down endogenous HeLa DYNC1H1 (hDHC), we verified that our observations were not an artefact of expression of mDHC-GFP in this background (Fig. S2a-h). In cells with no discernible mDHC-GFP, knockdown of hDHC led to dispersion of the Golgi marker GalT, while cells expressing low levels of mDHC-GFP (similar to those chosen for single-molecule imaging) had no effect on Golgi dispersion, showing that mDHC-GFP in these cells functions as expected (Fig. S2a). We verified hDHC knockdown using Western blot (Fig. S2b) and performed single-molecule analysis of dynein in control (Fig. S2c) or siRNA (Fig. S2d) treated hDHC-GFP HeLa cells. We verified the knockdown of hDHC by Western blot (Fig. S2e) and observed no difference in mDHC-GFP behavior viz. proportions of stationary, plus end-, and minus end-directed movement (Fig. S2f), residence time on MTs (Fig. S2g) and velocities (Fig. S2h). Further, the Western blots following hDHC depletion (Fig. S2b, e) indicated that the cells expressed mDHC-GFP only to a small extent over endogenous hDHC levels, since the DHC antibody used in these blots recognised both hDHC and mDHC. Taken together, mDHC-GFP behaves identically in the presence or absence of hDHC, functionally rescues hDHC depletion, and does not induce aberrant dynein behaviour and thus represents a good model to test dynein function *in cellulo*. To the best of our knowledge, these are the first observations of single molecules of dynein in mammalian cells and indicate that dynein likely exists in an inactive state inside the cell, similar to reports from *in vitro* studies (Splinter *et al*, 2012; McKenney *et al*, 2014; Schlager *et al*, 2014; Zhang *et al*, 2017).

### Dynactin remains persistently associated with MTs

Next, we aimed to visualise the dynamics of dynactin, the second player in the tripartite complex. Dynactin was first identified as a complex that was required for dynein-driven motility of vesicles *in vitro* (Gill *et al*, 1991). Several recent pieces of research have identified dynactin as an essential part of the active dynein complex (King & Schroer, 2000; Chowdhury *et al*, 2015; Urnavicius *et al*, 2015). Dynactin is a multi-subunit complex which binds to MTs independently of dynein via its N-terminal p150 subunit (Schroer, 2004). However, dynactin interacts poorly with dynein in the However, dynactin interacts with dynein poorly in the absence of cargo adaptor (Splinter *et al*, 2012; McKenney *et al*, 2014; Schlager *et al*, 2014).

The +TIP EB1 has been found to recruit another +TIP, CLIP-170, which in turn binds and clusters dynactin via its p150 subunit at growing MT plus ends (Watson & Stephens, 2006). MT plus ends thus decorated with dynactin also accumulated dynein at these sites, and evidence suggested that cargo transport was initiated when these MT plus ends contacted intracellular cargo (Vaughan *et al*, 2002; Moughamian *et al*, 2013). However, MT plus end-mediated initiation of dynein-driven transport appears to vary with cell type and context (Watson & Stephens, 2006; Kim *et al*, 2007; Tirumala & Ananthanarayanan, 2020). Therefore, using SD microscopy in combination with super-resolution radial fluctuations (SRRF, (Gustafsson *et al*, 2016)) we first quantified the localisation of p150 (Fig. 2a). Our high-resolution images revealed that only ~17% of the MT plus ends were enriched with p150, and p150 appeared bound along the entire length of the MT lattice (Fig. 2b). Further, by visualising hDHC along with EB1 using immunofluorescence (Fig. S2i-k), and quantifying the intensities of mDHC-GFP expressed in our cells, we concluded that the significant MT plus end localisation of dynein reported in earlier studies (Splinter *et al*, 2012; Kobayashi & Murayama, 2009) may represent an artefact of dynein overexpression (Fig. S3a, b). The dynactin complex, however, has been observed to show no MT-binding in the absence of dynein in *in vitro* assays (McKenney *et al*, 2014). To ascertain that the p150 spots we observed in these cells represented the entire dynactin complex, we used SRRF to visualise p150 in concert with another dynactin subunit, p62 (Fig. 2c). The p62 subunit of dynactin is located in the pointed end complex of dynactin (Schroer, 2004), and colocalisation of p62 with p150 would indicate the presence of the complete dynactin complex. We observed that 49±12 % (mean±s.d.) of the p150 spots colocalised with p62 (n=21,934/44,306 spots from N=2 independent experiments with 59 cells, Fig. 2d). Additionally, we used SRRF to visualise the localisation of p62 on MTs. The presence of p62 on MTs would indicate association with the MT of a subunit which does not normally do so unless it is part of the entire dynactin complex. Therefore, occurrence of p62 on the MT would imply localisation of the entire complex on the MT via p150 or dynein. We observed that 74±18% (mean±s.d.) of the p62 spots (n=59,715/79,639 spots from 1 experiment with 25 cells) were present on MTs (Fig. 2e, f). To corroborate our results using another super-resolution technique and to ensure our observations were not due to an image processing artifact, we utilised Airyscan confocal microscopy. First, we confirmed specificity of our p150 and p62 antibodies, showing they readily detect p150 and p62 expression constructs (Fig. S3c). We observed similar results to SRRF using Airyscan confocal microscopy with the p150 and p62 antibodies (Fig. S3d, e). Therefore, the complete dynactin complex is likely present along the entire length of the MT lattice, including the plus tip. Further, we observed that depletion of p150 through siRNA mediated silencing reduced the levels of both p150 and p62 along the MT lattice (Fig. S3f-k). To test that the perturbation of dynactin localisation along the MT lattice resulted in reduced dynein activity, we depleted p150 using siRNA mediated silencing and observed dynein behaviour. As expected, we observed a reduction in the proportion of dynein molecules that moved towards the MT minus ends with a concomitant increase in the proportion that moved towards the plus ends following p150 depletion (Fig. S4a-d). Taken together, our results indicate dynactin’s MT-binding function is likely essential for dynein’s activity *in cellulo*.

**Fig. 2.**
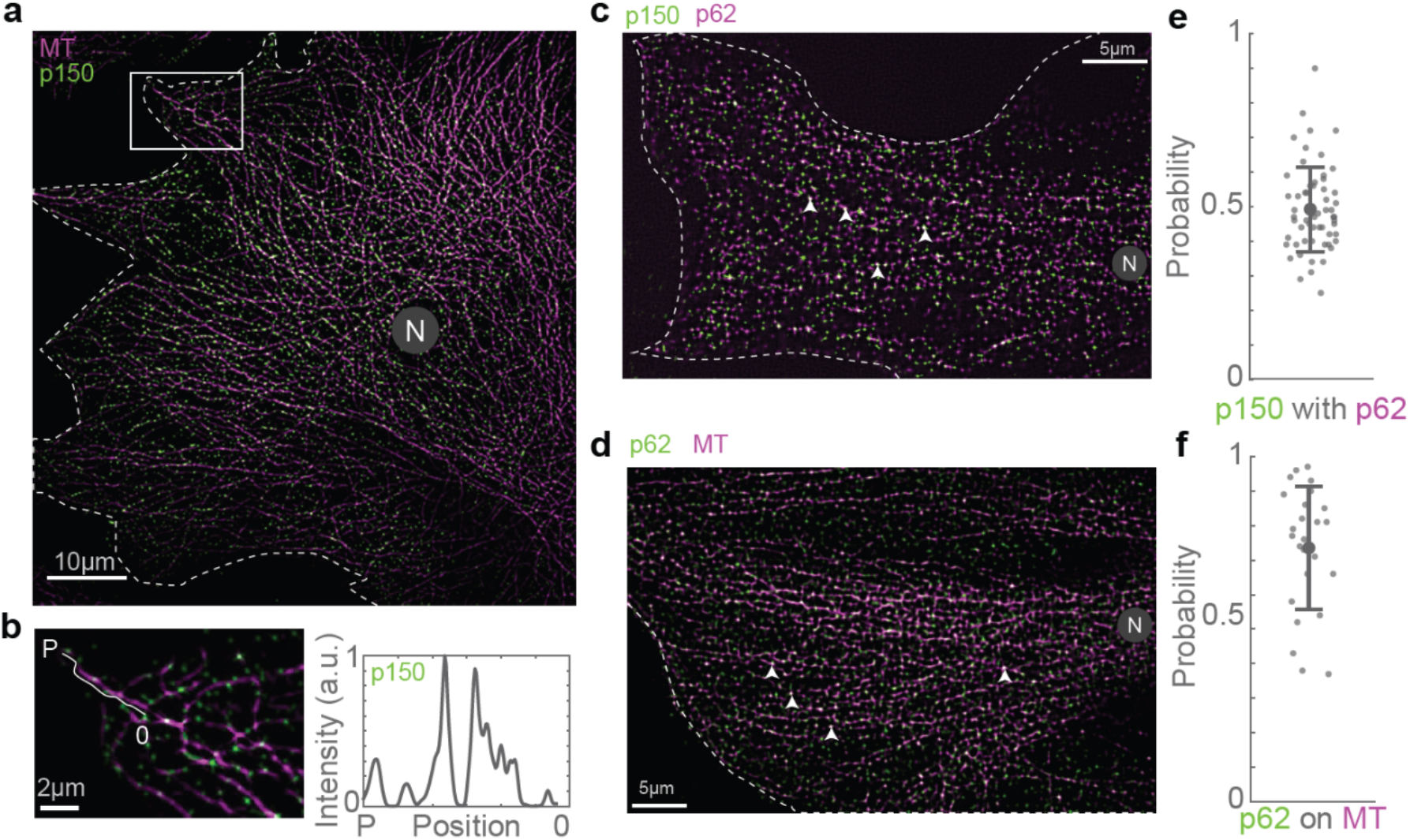
The dynactin complex binds along the entire length of the MT. **a**, Immunofluorescence image of MT (magenta) and p150 (green) obtained using SD microscopy + SRRF. **b**, Enlarged view of the area marked with the white rectangle in **a** and the line profile of p150 intensity along the length of a representative MT from the plus end (‘P’) to −6 *μm* from the plus end of the MT (O’). **c**, Immunofluorescence image of p62 (green) and p150 (magenta) obtained using SD microscopy + SRRF. The white arrowheads indicate representative p150 spots that also contain p62. **d**, Immunofluorescence image of MT (magenta) and p62 (green) obtained using SD microscopy + SRRF. The white arrowheads indicate representative p62 spots that occur on the MT. **e**, Histogram of the probability of cooccurrence of p62 with p150, indicating a high likelihood of presence of the entire complex at a p150 spot. **f**, Histogram of the probability of cooccurrence of p62 on the MT, which points to a high likelihood for the presence of the entire dynactin complex on the MT. In **a**, **c** and **d**, ‘N’ marks the location/direction of the nucleus.

### Endosomes remain close to MTs and move in short bursts

We next sought to understand how dynein interacted with the third component of the active complex - the cargo adaptors. The endosomal cargo adaptors, Hook and BicD2 proteins, have been observed to remain persistently bound to their respective cargo (Bielska *et al*, 2014; Matanis *et al*, 2002). Therefore, we used endosomal cargo as a proxy for the cargo adaptor, which confers the additional benefit of visualising the movement of the entire cargo and adaptor complex. To avoid artefacts from overexpressing fluorescently tagged Rab5 to visualise early endosomes (Nielsen *et al*, 1999), we employed cells that had taken up Alexa^647^ conjugated 10kDa dextran or EGF (Fig. S5a).

Small molecular weight dextran enters the cell through all active endocytic mechanisms via fluid phase endocytosis, and uptake is therefore relatively independent of receptor-mediated endocytosis (Li *et al*, 2015; Rennick *et al*, 2021). On the other hand, EGF is a ligand that binds to EGF receptor on the cell membrane, which at the concentrations used in this study, is taken up by clathrin-mediated endocytosis (Goh *et al*, 2010). While both these cargoes eventually end up at lysosomes near the minus ends of MTs, the timescales of movement of these cargoes is vastly different, with dextran requiring hours (Humphries *et al*, 2011) to reach lysosomes and EGF doing so in tens of minutes (Futter *et al*, 1996). We sought to understand how dynein’s short run times act to ensure cargoes reach the lysosomes in different time scales.

Following a short pulse and chase, we observed dextran in Rab5-positive compartments (Fig. S5b, Movie S3). EGF has also been shown to be in Rab5-positive compartments within the timescales of our chase observations (Leonard *et al*, 2008), indicating that both dextran and EGF were in early endosomes in our experiments. We probed the localisation of dextran and EGF vesicles with respect to the MT. First, we imaged dextran vesicles and MTs in live cells and observed that the vesicles were in proximity to the MTs (Fig. 3a). We also observed that the dextran vesicles remained close to MTs even while they had no observable tether to the MTs via motor proteins and were therefore stationary (Movie S4) in our minute-long live cell timelapse images. We then used SRRF to visualise Rab5 in concert with p62 and observed that ~70% of the Rab5 spots colocalised with p62 (n=8,795/12,848 vesicles from N=2 independent experiments with >20 cells each), indicating that cargo and dynactin are in proximity to each other on the MT (Fig. 3b, c). Finally, we employed CLEM to visualise the location of dextran and EGF endosomes with respect to MTs. We observed that in both instances, the endosomes were along MTs, indicating that endosomal cargo remained within ~ 30 nm of MTs (dextran: 31±15 nm, from 23 endosomes, N=5 cells; EGF: 26±18 nm, from 33 endosomes, N=6 cells), likely along with dynactin (Fig. 3d-j). Interestingly, the size of the dynactin complex is ~35 nm (Hodgkinson *et al*, 2005), comparable to the distances observed between dextran and EGF endosomes from MTs, indicating dynactin may hold endosomes to MTs.

**Fig. 3.**
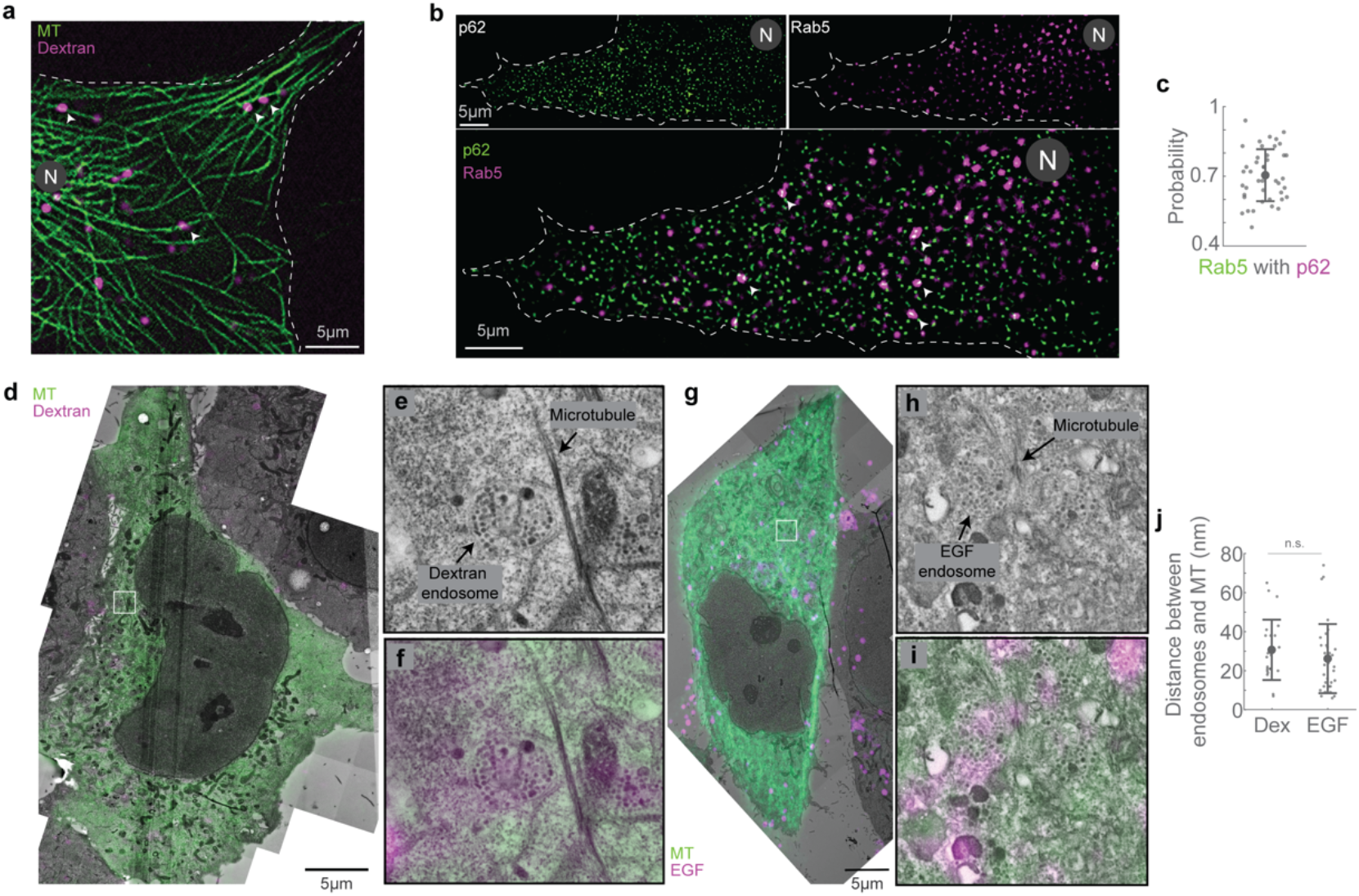
Endosomes remain close to MTs. **a**, SD + SRRF image of MTs (green) and dextran vesicles (magenta) in live cells. The white arrowheads indicate representative vesicles on the MT. **b**, SD + SRRF image of p62 (top left), dextran vesicles (top right) in live cells and their merge (bottom, p62 in green and dextran in magenta). The white arrowheads in the merged image indicate representative vesicles that colocalise with p62 and are shown as green and magenta arrowheads in the p62 and dextran images respectively. **c**, Histogram of the probability of cooccurrence of p62 with Rab5, indicating a high likelihood of dynactin being present in a complex with endosomal cargo. **d**, Overlay of confocal images of MT (green) and dextran vesicles (magenta), and EM images of the same cell (grey). **e**, EM image of the region indicated with the white square in **d**, showing a representative MT and dextran endosome. **f**, Confocal fluorescence image of MTs (green) and dextran (magenta) of the region in **e**. **g**, Overlay of confocal images of MT (green) and EGF vesicles (magenta), and EM images of the same cell (grey). **h**, EM image of the region indicated with the white square in **g**, showing a representative MT and EGF endosome. **i**, Confocal fluorescence image of MTs (green) and EGF (magenta) of the region in **h**. In **a** and **b**, ‘N’ marks the location/direction of the nucleus. **j,** Quantification of the measured distance between dextran (‘Dex’) and EGF endosomes and MTs. ‘n.s.’ indicates no significant difference (p=0.3), one-way ANOVA, Tukey Kramer post-hoc test.

To understand if endosomal vesicles could diffuse away from their original locations on the MT upon motor unbinding, we depolymerised MTs using nocodazole and visualised the subsequent movement of the vesicles (Fig. S5c, Movie S5). Confirming previous findings on Rab5-positive compartments (Flores-Rodriguez *et al*, 2011; Zajac *et al*, 2013), we measured a lower diffusion coefficient for the vesicles in the absence of MTs (Fig. S5d), likely implicating the role of high intracellular crowding in constraining vesicle diffusion, also previously noted by others (Zajac *et al*, 2013; Weiss *et al*, 2004; Szymanski & Weiss, 2009; Ernst *et al*, 2012; Sokolov, 2012). This intracellular crowding may also obviate the need for dynactin to act as a tether between endosomes and MTs.

We next visualised dextran and EGF vesicles in 100s-timelapses and observed that the directed runs were sparse (Fig. 4a, b, Movie S6, S7), with only ~33% of dextran vesicles (65/196) and ~43% EGF vesicles (92/214) moving >1 μm during this time (Fig. 4c). Strikingly, both the dextran and EGF vesicles displayed uninterrupted minus end-directed runs that lasted only ~ 0.6 s on an average (Fig. 4d) (0.6±0.2 s (mean±s.d.), n=196 and 214 for dextran and EGF vesicles respectively from N=2 independent experiments). Interestingly, the time between two consecutive runs, the pause time, reflected the time scales of minus end-directed movement of EGF and dextran, with EGF vesicles having an average pause time that was 30% shorter than that of dextran vesicles (0.4 s vs. 0.6 s, Fig. 4e). These results are comparable to the run- and-pause behaviour observed for endosomal cargo in previous studies (Flores-Rodriguez *et al*, 2011; Zajac *et al*, 2013).

**Fig. 4.**
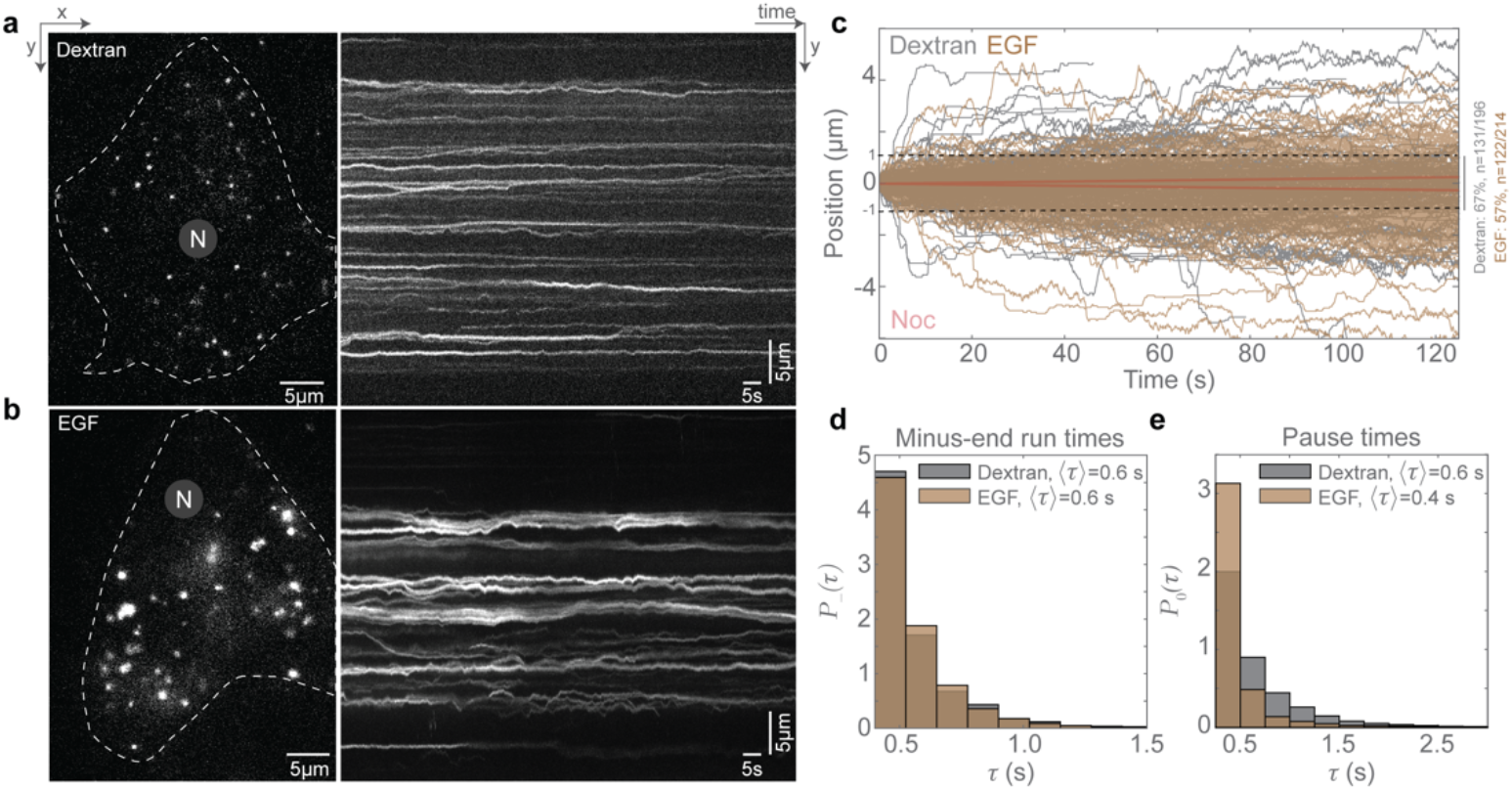
Dextran and EGF vesicles display distinct movement kinetics. **a**, HILO image from a time-lapse of dextran vesicles in HeLa cells (left) and the ∞rresponding kymograph (right), **b**, HILO image from a time-lapse of EGF vesicles in HeLa cells (left) and the corresponding kymograph (right), **c**, Plot of position versus time of dextran vesicles (grey) and EGF (brown) in HeLa cells, and the predicted movement of endosomes after nocodazole treatment in the absence of MTs (red). The dashed black lines indicate absolute movement of 1 μm, with 67% of dextran vesicles and 57% of EGF vesicles moving less than 1 μm during the time-lapse imaging. **d**, Probability distribution *P*-(*r*) of the minus end-directed run times for dextran (grey) and EGF (brown) vesicles, **e**, Probability distribution *P*_O_(*τ*) of the pause times for dextran (grey) and EGF (brown) vesicles. In **a** and **b**, ‘N’ marks the location/di recti on of the nucleus. In **d** and **e**, the average run/pause time (〈*τ*〉) for dextran and EGF vesicles are indicated.

### Transient dynein attachment to cargo complexes is sufficient to drive typical cargo trafficking

Thus far, we have discovered that dynein transiently interacts with the MTs while dynactin-cargo complexes are maintained close to MTs. We therefore sought to understand how dynein interacted with the dynactin-cargo complexes on MTs. First, we performed dual-colour imaging of dynein and endosomal vesicles, and observed that vesicles that had dynein signal were, as expected, more likely to move towards the minus end of MTs (Fig. S5e-g). Next, by comparing the intensity of mDHC-GFP on endosomal vesicles to single molecule binding events in cells that were depleted of hDHC, we estimated that on average there were 1-2 dynein molecules bound to a vesicle (n=44 endosomes, from N=2 independent experiments with >25 cells, Fig. S5h, i). Finally, to verify if single molecules of dynein could be activated and perform minus end-directed movement when they stochastically bound to MTs and encountered cargo complexes, we performed fast dual-colour HILO imaging of single dynein molecules, along with dextran and EGF vesicles. In these videos (Movies S8, S9), we observed instances where previously stationary dextran and EGF vesicles started moving together with mDHC-GFP towards the MT minus ends upon dynein binding (Fig. 5a-d, n=8/9 and n=3/3 such events for dextran and EGF respectively from N=2 independent experiments with at least 15 cells). We note that the examples in Movies S8 and S9 are rarer, longer events, and we additionally observed shorter events of dynein binding and effecting dextran and EGF vesicle movements (Fig. S5j, k, n=19 and n=14 short events for dextran and EGF respectively from N=2 independent experiments with at least 15 cells).

**Fig. 5.**
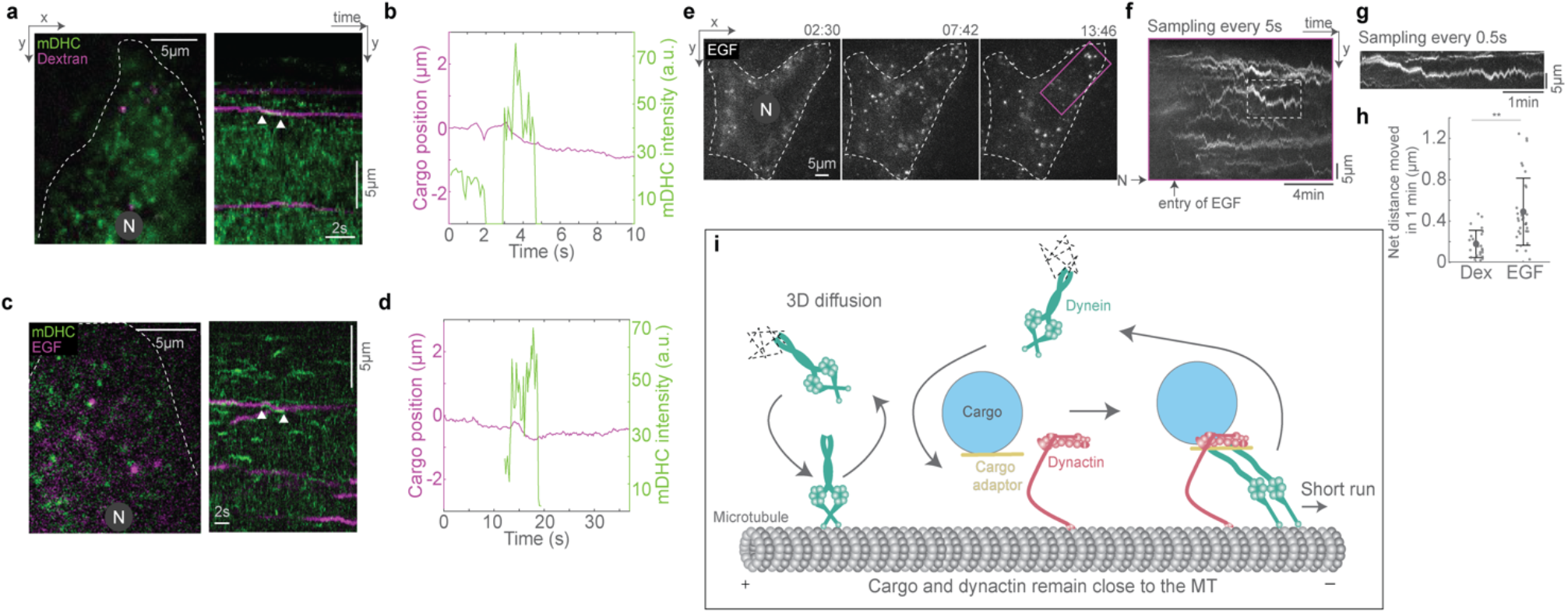
Cargo capture and dynein activation are coupled. **a**, Image from the first frame of a time-lapse video (left) of dynein (green) and dextran (magenta), and the corresponding kymograph (right). The white arrow heads point to dynein and dextran vesicle moving together towards the minus end. **b**, Plot of position vs time of the dextran vesicle (magenta) indicated in **a**, alongside the intensity of dynein on that vesicle (green), showing a short minus-end directed run of the vesicle upon dynein-binding. **c**, Image from the first frame of a time-lapse video (left) of dynein (green) and EGF (magenta), and the corresponding kymograph (right). The white arrow heads point to dynein and EGF vesicle moving together towards the minus end. **d**, Plot of position vs time of the EGF vesicle (magenta) indicated in **c**, alongside the intensity of dynein on that vesicle (green), showing a short minus-end directed run of the vesicle upon dynein-binding. **e**, Images from time-lapse video of EGF entry and subsequent movement within cells. Time is indicated above the images in mm:ss. **f**, Kymograph of the region indicated by the magenta rectangle in **e**, with the timelapse images sampled every 5 s showing entry of EGF and the net movement of endosomes towards the nucleus. **g**, Kymograph of the trajectory marked by the white dashed rectangle in **f**, where the timelapse images were sampled every 0.5 s. In **a**, **c**, and **e**, ‘N’ marks the location/direction of the nucleus. **h**, Quantification of the net distance moved by dextran (‘Dex’) and EGF endosomes in 1 min. Asterisks indicate significant difference, p=4×10^-5^, one-way ANOVA, Tukey Kramer post-hoc test. **i**, Schematic of dynein’s cargo search mechanism: stochastic binding of dynein to the MT at a location proximal to the cargo-adaptor complex leads to a short minus end directed run, which terminates upon the unbinding of dynein.

We next queried whether the transient movements driven by dynein were sufficient to drive EGF and dextran vesicles to perinuclear regions of the cell in timeframes consistent with their delivery to lysosomes. We visualized the uptake and subsequent movement of EGF endosomes over a 17-min period using HILO microscopy. We observed robust movement of the EGF endosomes in the net minus end direction during the duration of imaging (Fig. 5e, f, Movie S10). Strikingly, movements that appeared as long-range at a low sampling rate (Fig. 5f), showed intrinsic short runs when sampled at a higher rate (Fig. 5g), confirming the short range movements such as those observed Fig. 4 are cumulatively sufficient to drive expected cargo transport.

We measured the net movement of dextran and EGF endosomes in these longer time-lapses and observed that these endosomes moved a net distance of 0.2±0.1 *μm* and 0.5±0.3 *μ*m (mean±s.d., from n=24 dextran endosomes from 12 cells, N=2, and n=32 EGF endosomes from 9 cells, N=2) in 1 min respectively (Fig 5h). Considering the average distance between the cell periphery and a lysosome to be 10 *μ*m, a dextran endosome in our experiments would require 50 min and an EGF endosome, 20 min on an average to reach the lysosome, which are in agreement with previously measured time scales of movement of these cargoes (Humphries *et al*, 2011; Futter *et al*, 1996). To confirm that our experimental conditions were optimised for observing these expected trafficking timeframes, we performed additional quantification of the distance of dextran- and EGF-containing endosomes from the cell periphery over time (Fig. S6a-f), as well as their co-occurrence with the late endosomal marker Rab7 and the lysosomal marker LAMP1. These investigations revealed that the behaviour of these endosomes was in agreement with previous literature, wherein EGF endosomes occurred at larger distances from the periphery, and also were more likely to be Rab7 and LAMP1 than dextran-containing endosomes at the same time point after internalisation (Fig. S6g, h), with EGF endosomes travelling a net minimum distance of 0.25 μm per minute and dextran endosomes 0.2 μm per minute (Fig. S6i). Altogether, the higher temporal resolution of our experiments enabled us to discern the dynein-driven stop-and-go behaviour of endosomal cargo that is ultimately sufficient to drive movement to its intracellular destination in time frames consistent with those previously observed.

### Conceptual model of dynein’s cargo attachment and subsequent movement within cells

Taken together, we observed that: (i) single molecules of dynein bind and unbind stochastically with MTs (Fig. 1b), (ii) if a dynactin-cargo complex is found close to this attachment location of dynein to the MT, a minus end-directed run of the dynactin-cargo-motor complex is effected (Fig. 5a-d), (iii) the detachment of dynein from the cargo concludes this run (Fig. 5i), (iv) the detached dynein motor is free to diffuse back into the cytoplasm, whereas dynactin and the cargo remain paused and remain close to the MT (Fig. 3), (v) the long-range movement of endosomal cargo requires the repeated binding-unbinding of dynein to the dynactin-cargo complex, (vi) the resulting motion of the cargo consists of short minus (and plus) end-directed runs punctuated by long pauses (Fig. 4c). Our experimental techniques - HILO, SD, SRRF, Airyscan and CLEM - provided high spatiotemporal data for the dynamics of motors and dynactin-cargo complexes. In fact, this not only allowed us to quantify the stochastic kinetics of individual dynein motors to extract its detachment rates (Fig. 1f), but also allowed us to explore the long-time dynamics of cargo with high precision (Fig. 4c, 5e-g). As such, we sought to quantify the statistical dynamics of endosomal cargo by measuring the distributions of the cargo residence time of both dextran and EGF in the moving and paused states. Before discussing these results in detail (Fig. 7–9), we develop stochastic models at two levels of descriptions that can quantitatively account for these distributions of cargo dynamics.

### A stochastic model captures the kinetics of motor-driven cargo movements

To study the stochastic dynamics of motors and cargo units capable of activating the motor (which include dynactin and adaptor for dynein, henceforth referred to as ‘cargo complexes’), we modelled the MT as a discrete one-dimensional lattice of *N* binding-sites each of size Δ*x* (=0.05 *μ*m) (Fig. 6a). Our model included both dyneins and kinesins interacting with the MT and cargo complexes. At each site *i* of the MT, exchange kinetics with a large cytoplasmic reservoir caused dynein (kinesin) motors to bind at a Poisson rate 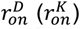 to the MT and unbind from the MT at a Poisson rate 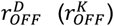 to the cytoplasm. Note that this binding-unbinding process was stochastic and occurred uniformly throughout the length of the MT. If a cargo complex was found at the binding location of a motor protein, then the motor-cargo complex translocated along the MT with a speed *v^D^* towards the minus end in the case of dyneins and with a speed *v^K^* towards the plus end in the case of kinesins. Dynein (kinesin) motors detached from the MT after a mean residence time 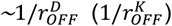, which also ended the motion of the cargo complex. From this point on, the cargo complex remained stationary, but still in proximity to the MT. In other words, we completely neglected the 3D diffusion of the cargo-adaptor complexes since our data suggested that cargo diffusion is negligible (Fig. S5d). Furthermore, we only considered a single cargo complex at a time and disregarded any steric interactions between them. The chemical master equation describing this Markovian stochastic process is described in detail in the Supplementary Text. When we restricted the number of motors that could be bound to each site of the MT (*M*) to one, this stochastic motor-cargo complex dynamics model (referred to as the ‘MCD model’ henceforth) generated cargo trajectories that underwent pauses, minus end-directed and plus end-directed runs (Fig. S7). However, in this instance, the only transitions that occurred were between paused and moving (plus or minus) states of the cargo complex, which were not truly reflective of our experimental observations.

**Fig. 6.**
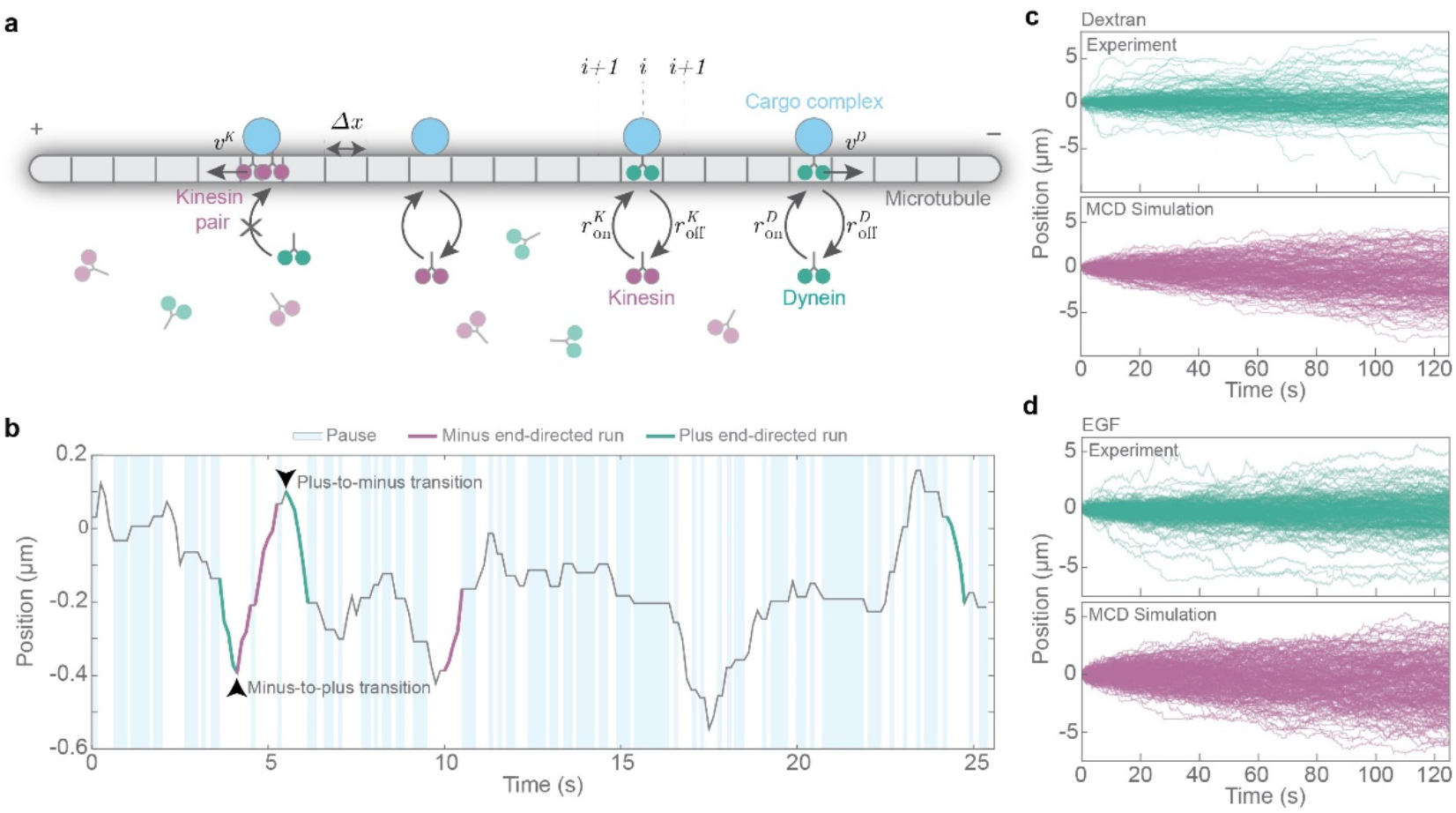
MCD model for dynein’s cargo capture and activation a, Schematic of the MCD model depicting the MT (grey), with lattice points separated by a distance. Cargo complexes (blue) are positioned randomly along the MT and do not diffuse away. Dynein (green) and kinesin (magenta) motors bind and unbind stochastically to the **MT lattice points with rates** 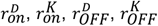, **respectively. If a cargo-complex is present at the site of motor binding** (*i*), **the motor could hop to the adjacent site** *i* + 1 or *i* – 1, with a rate *v^K^* or *v^D^* for kinesin and dynein respectively. Note that we if two motors are already present at a lattice point (e.g.: ‘Kinesin pair’), no more motors can bind to the same site. We used periodic boundary conditions to simulate this model on an MT of length *N* **Δ** *x*. **b**, An example position vs. time trajectory of a dextran vesicle obtained from experiment (grey line), showing representative pauses (blue rectangles), minus end-directed runs (solid magenta lines), plus end-directed runs (solid green lines), and transitions from minus-to-plus and plus-to-minus end-directed runs (black arrowheads). **c**, Position vs. time trajectories of dextran vesicles from experimental data (green, top) and from the MCD simulation (magenta, bottom). **d**, Position vs. time trajectories of EGF vesicles from experimental data (green, top) and from the MCD simulation (magenta, bottom). Note that the experimental data in **c** and **d** are identical to those in Fig. 4c and have been reproduced here for comparison.

Typical experimental trajectories of the cargo showed discrete paused and moving states (Fig. 6b for dextran). In addition to the transitions between pauses and runs, we observed direct transitions between minus end-directed runs and plus end-directed runs. This, however, is not possible unless the cargo complex was moved by at least a pair of motors consisting of one dynein and one kinesin. In other words, our experimental data (Fig. 6b) indicated that there were antagonistic pairs of motor proteins bound to the cargo complex (see Fig. S8 for the output of the MCD model with M=1). In fact, our experiments with dual imaging data of single molecules of dynein and cargo indicated that there were one to two dyneins bound to the cargo (Fig. S5h). It was thus reasonable to assume that we would also have pairs of kinesins, and dynein-kinesin pairs bound to a cargo complex. Note that cargo translocation could be affected by both single motors as well as pairs of motors.

We therefore extended our MCD model to the case where a maximum of *M*=2 motors could be bound to each site of the MT. Note that even with *M*=2, we assumed that the motors hopped at the same speeds, *i.e*., at rates *v^D^*/Δ*x* and *v^K^*/Δ*x*, across the lattice sites of the MT. We also assumed that the antagonistic pair could move in either direction at the appropriate rates. Further, we neglected any cooperative binding between multiple motors. We discuss in detail the chemical master equation associated with this model in the Supplementary Text, and also its stochastic simulation via the Gillespie algorithm (Gillespie, 1977; Erban & Chapman, 2019). With values for the parameters 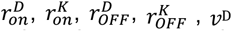 and *v^K^* that compared reasonably well with their estimates from experiments (see table in Fig. 9c), the MCD model generated cargo trajectories that were comparable to empirical results (Fig. 6c and d). However, taking advantage of our long-term and high temporal resolution imaging data, we next asked whether the model could quantitatively account for the residence time distributions of the cargo in the various states, and if there was a simplified description for the dynamics of the cargo complexes that themselves behaved as “super-motors”, displaying motion which was markedly different from simple Brownian motion on the MT.

### The emergent dynamics of cargo is captured by a 3-state run-and-tumble model

Typical trajectories, such as the one shown in Fig. 6b, revealed short periods of directed motion of the cargo complexes interspersed with frequent pause events. This kind of stochastic motion was *not* characteristic of the simple, and well-known, thermal Brownian motion. Rather, this movement was reminiscent of the classic run-and-tumble motion wherein bacteria run in almost straight-line paths for a certain run-duration after which they tumble while being stationary for a tumble-duration, and then run again in a randomly chosen direction (Berg *et al*, 2004). Statistical physics models of such run-and-tumble particles (RTP) have revealed very interesting features of these persistent and active random walks (Tailleur & Cates, 2008; Bechinger *et al*, 2016; Malakar *et al*, 2018). However, in these well-studied RTP models, the tumble is, typically considered to be an instantaneous event. Several studies have explored models, with three distinct states–two moving states (with movements in opposite directions), and a paused state– for cargo that arise from underlying motor movement (Smith & Simmons, 2001; Bressloff & Newby, 2013).

Our MCD model discussed in the previous section led to an emergent picture for the dynamics of the cargo complexes as RTPs moving on a one-dimensional MT with three distinct states: *minus end-directed, paused* and *plus end-directed*. In more detail, these states resulted from the following situations (i) a minus-end directed run of the cargo complex occurred where there were motor configurations dominated by dyneins, (ii) cargo complexes were paused when there were no motors (of either kind) within a proximal region on the MT, and (iii) a plus-end directed run occurred when kinesins were the primary movers of the cargo complexes. As such, the effective movement of the cargo was captured by three discrete states with distinct transition rates between them.

We captured the dynamics of the cargo complexes discussed above in a 3-state RTP model. Specifically, as depicted in Figure 7a, stationary cargo could transition to the plus (minus) end-directed state at a rate *γ*_0p_(*γ*_m_), while moving cargo could transition to the stationary state at rates *γ*_m0_ (from the minus end-directed state) and *γ*_p0_ (from the plus end-directed state). Further, we considered direct transitions between the plus and minus end-directed states at rates *γ*_pm_ and *γ*_mp_ respectively since these transitions were apparent in our experimental data (Fig. 6b). In the plus end-directed state, the cargo translocated with a velocity *u* = +*u^p^* and in the minus end-directed state, the translocation velocity was *u* = -*u^m^*. In the stationary state, by definition, the cargo was immobile, i.e., *u* = 0. Therefore, denoting 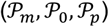 as the probabilities of finding the cargo in the minus end-directed, stationary and plus end-directed states respectively, the master equations governing their Markovian time-evolution were:

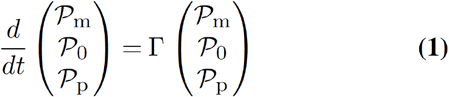

where the transitional matrix *Γ* was:

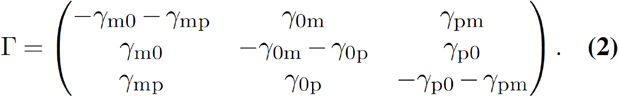

**Fig. 7.**
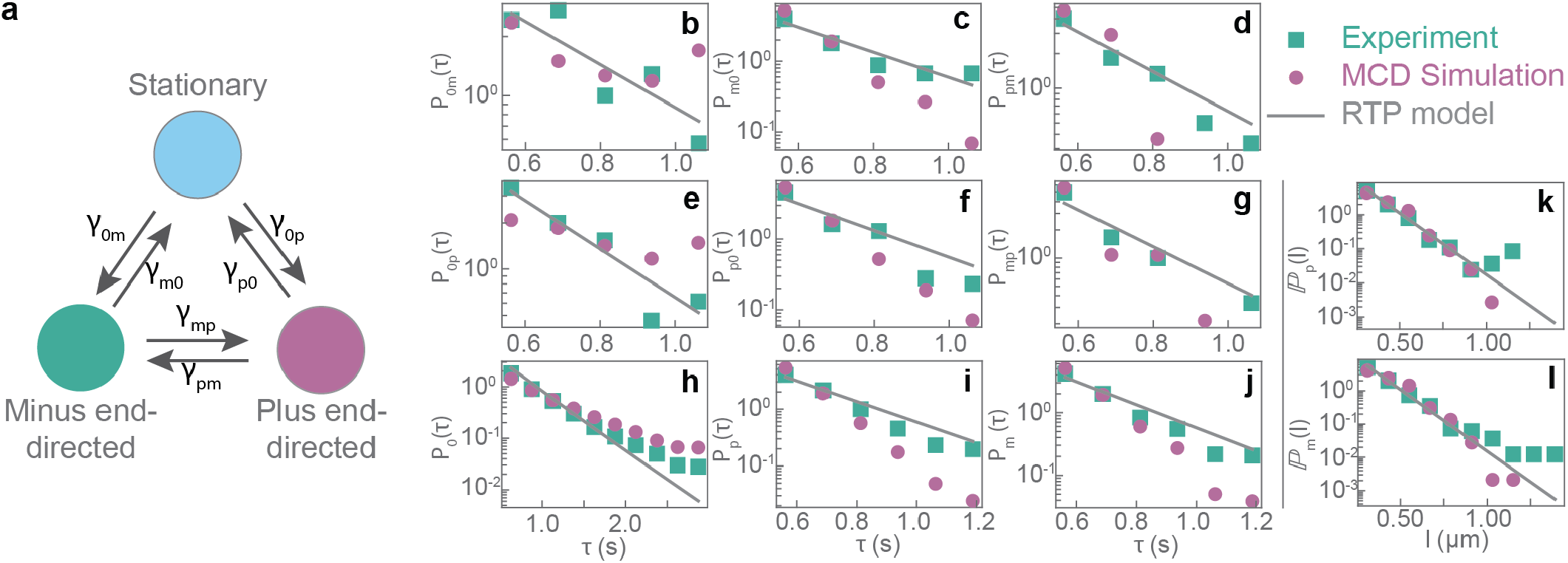
The 3-state RTP model and its fits for the movement of dextran vesicles. **a**, Schematic of the RTP model depicting the 3 states: stationary, minus end-directed and plus end-directed, and the transition rates between them, i.e. *γ*_0m_, *γ*_m0_-rates of transition from the stationary to minus end-directed state and vice versa; *γ*_0p_, *γ*_p0_-rate of transition from the stationary to the plus end-directed state and vice versa; *γ*_mp_, *γ*_pm_-rates of transition from the minus end-directed to the plus end-directed state and vice versa. In **b-g**, experimental data (green squares) are compared with the output from MCD simulations (magenta circles) and the predictions of the RTP model (grey lines) for the probability distributions of the cargo waiting time (*τ*) while transitioning from the stationary to the minus end-directed state (**b**), minus end-directed to stationary (**c**), plus to minus end-directed (**d**), stationary to plus end-directed (**e**), plus end-directed to stationary (**f**), and minus end-directed to plus end-directed (**g**). Further, the probability distributions of the pause-times (**h**), the run times in the plus end-directed state (**i**), and the run times in the minus end-directed state (**j**) are shown in **h-j**, while **k** and **l** show the distributions of the the run lengths (*l*) in the plus end-directed and minus end-directed states respectively. The solid grey lines represent fits to the experimental data from the RTP model. The MCD model results allowed pairs of motors to transport cargo-complexes, i.e., *M*=2. Note that he plots (**b-l**) are in log-linear scale.

Note that in these emergent discrete states, the cargo complexes moved with the appropriate velocity. In other words, the position *x* of the cargo complex changed with time according to *dx/dt* = *u*, where the velocity *u* could take either of the three possibilities {−*u*^m^, 0, *u*p}.

The stochastic dynamics captured by equation Eq. (1) predicts that the distribution *P*_ab_(*τ*) of the residence time (equivalently waiting time) in the *a*-state before jumping to the *b*-state is an exponential distribution with the rate constant *γ*_ab_ (here a, b could take either of the values m, 0 or p). For instance, the distribution of the residence times of the cargo complex in the paused state before transitioning to the minus end-directed state would be 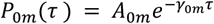, where *A*_0*m*_ is an amplitude factor. Similar considerations applied to the waiting time distribution for the other transitions.

We next asked whether our high-temporal resolution, long-term data for the positions of the cargo complexes was consistent with the waiting time distributions obtained from this coarse-grained 3-state RTP model. To this end, we partitioned the trajectory of the cargo complexes into discrete states (Fig. 6b, see SI Methods), measured the waiting time for each transition depicted in Fig. 7a, and fitted exponential functions to the probability density functions of the residence times *τ*. We found a good match between our empirical data and the distributions *P*_ab_(*τ*) as shown in Fig. 7(b-g) for the case of dextran vesicles. We employed this procedure to extract the effective rate constants yab for a transition of the cargo complex from the a-state to the b-state (see the table in Fig. 9c). Notice that the cargo could exit the stationary/paused state via two independent pathways: transitioning into the minus end-directed state or transitioning into the plus end-directed state. As such, the distribution of the pause times was a mixed distribution 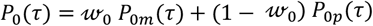 where 0 ≤ *w*_0_ ≤ 1 is a weight factor. To check whether this prediction worked, we fixed the rates *γ*_0*p*_ and *γ*_0*m*_ to the values obtained by the fits in Fig. 7b and Fig. 7e respectively, and fit the form of *P*_0_(*τ*) to the empirical data. The excellent fit seen in Fig. 7h confirms the validity of our 3-state RTP model. Similar considerations applied for the distribution 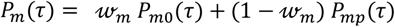 of minus end-directed run times (Fig. 7i), and the distribution 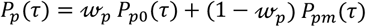 of plus end-directed run times (Fig. 7j). Further, in the minus (plus) end-directed states, we measured the distribution of the run lengths 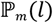 and 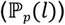. Since the cargo complexes only moved when they were in the *m*-or *p*-states, the run length distributions were expected to be related to the run time distributions. Specifically, we expected: 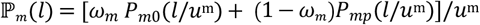, and 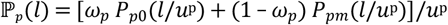. The excellent fits of the empirical data to these distributions seen in (Fig. 7j-k) allowed us to extract the emergent speeds *u*^m^ and *u*^p^ of the cargo complexes.

We obtained more compelling evidence to this emerging picture when we were able to match the results of the MCD simulation, depicted in Fig. 6a, for the various residence time and run length distributions of the empirical data and the RTP model. We repeated this procedure for the case of EGF vesicles and obtained similar agreement between the empirical data, the MCD simulation and the RTP model as shown in Fig. 8. The parameter values both in our MCD simulations and those obtained in the RTP model are available in Fig. 9c. The predictive power of our approach was revealed when we examined the dependence of the average run length 〈*l*〉 as a function of the run time *τ* in the case of both dextran and EGF. Since all parameters in the RTP model were known, we compared the empirical data and the MCD simulation data with the predictions 〈*l*〉 = *u*^p*τ*^ and 〈*l*〉 = -*u*^m^*τ* for plus end-directed and minus end-directed runs respectively. The remarkable agreement seen in Fig. 9a and b confirmed that our simulation and modelling approaches provide an accurate description of the experimental data. So too, the scaling of emergent cargo velocities with run time points to the participation of at least 2 motors in the cargo trafficking process. Indeed, the output from the MCD model with *M* = 1 did not match the experimental data (Fig. S8) or fit the RTP model (Figs. S8, S9a).

**Fig. 8.**
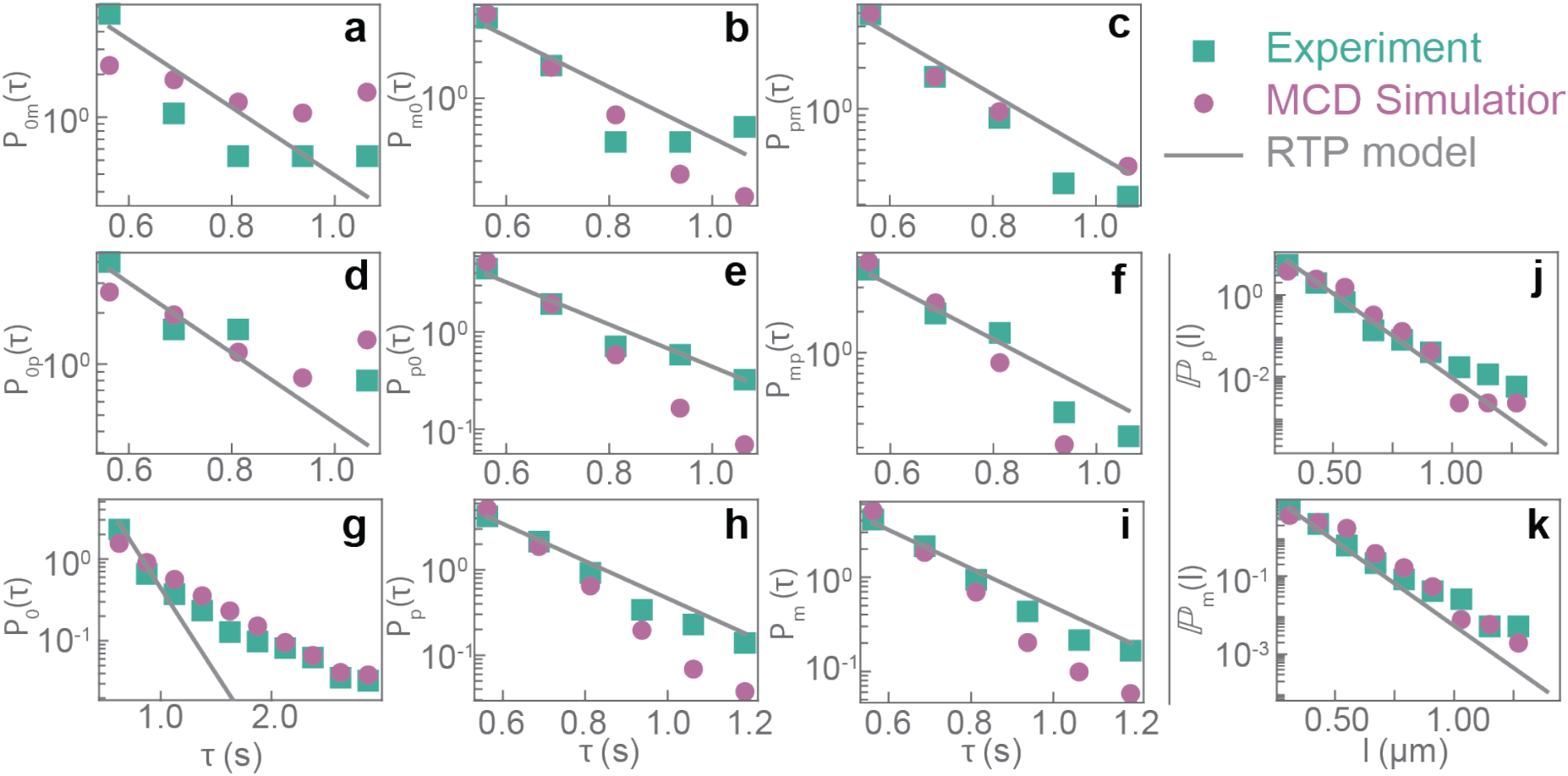
Fits of the 3-state RTP model for EGF vesicle trajectories. In **a-f**, experimental data (green squares) are compared with the output from MCD simulations (magenta circles) and the predictions of the RTP model (grey lines) for the probability distributions of the cargo waiting time (*τ*) while transitioning from the stationary to the minus end-directed state (**a**), minus end-directed to stationary (**b**), plus to minus end-directed (**c**), stationary to plus end-directed (**d**), plus end-directed to stationary (**e**), and minus end-directed to plus end-directed (**f**). Further, the probability distributions of the pause-times (**g**), the run times in the plus end-directed state (**h**), and the run times in the minus end-directed state (**i**) are shown in **g-i**, while **j** and **k** show the distributions of the run lengths (*Z*) in the plus end-directed and minus end-directed states respectively. The solid grey lines represent fits to the experimental data from the RTP model. The MCD model results allowed pairs of motors to transport cargo-complexes, i.e., M=2. Note that the plots **a-k** are in log-linear scale.

**Fig. 9.**
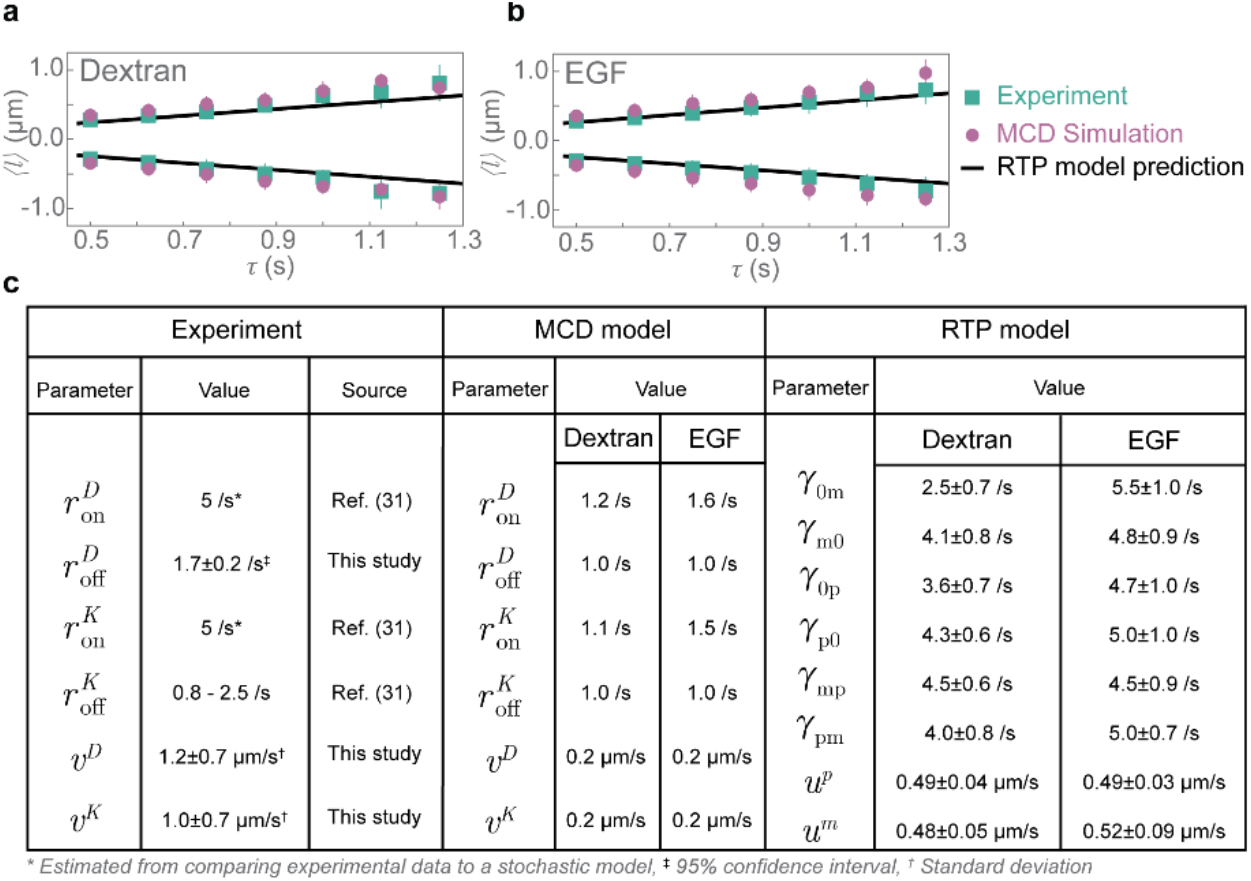
Run lengths of cargo scale with run time, Plot of average run length 〈*l*〉 as a function of run time *τ* for dextran vesicles (**a**) and EGF vesicles (**b**), with data from experiments (green squares) and output from the MCD model with *M* = 2 (magenta circles). The black solid line represents the prediction of the RTP model using the parameter listed in the table below. **c** Table containing the parameters that were measured for single dynein and kinesin molecules in this study and other experiments (‘Experiments’), those that were employed in the MCD model (‘Stochastic model’), and those for the cargo movement that obtained from fits to the experimental data from the RTP model (‘RTP model’) for both dextran and EGF.

More sophisticated analysis of cargo trajectories either Levy walks (Chen *et al*, 2015) or as those resulting from fractional Brownian motion (Han *et al*, 2020) could be employed in future to ask if the effective dynamics of the cargo (averaged over the dynamics of the motors) would fall into either of the above classes. It is thus apparent that our MCD model, which explicitly included the dynamics of motors, led to a coarse-grained emergent view wherein the dynamics of the cargo complexes were governed by a 3-state RTP model. However, an important point to note is that it is a non-trivial task to find analytical relations between the emergent parameters at the RTP level 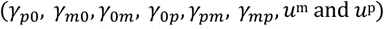 and the parameters in the model (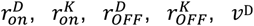 and *ν*^K^). Nevertheless, the parameters used in our simulations revealed that the on-rates for both dynein and kinesin are higher by ~ 30% for EGF as compared to dextran. Remarkably, our experimental data showed that EGF vesicles had an average pause time which was ~ 30% shorter than that of dextran vesicles (Fig. 4e). This difference in on-rates and thereby the pause times, could have arisen from an increased number of adaptors for dynein (and kinesin) on EGF-containing vesicles compared to dextran-containing vesicles and will be the subject of future studies. Together, we developed a framework for considering cargo movement in light of a 3-state RTP model which could be used to infer not only underlying motor kinetics, but also predict long-term cargo movement and fates, such as in differential cargo sorting.

## Discussion

In this work, we established a technique to visualise single molecules of dynein in living cells, and observed their interaction with the MT, and other components of the tripartite complex required for its activation. We discovered distinct localisation and kinetics of dynein, dynactin, and the cargo complex. While dynein interacted stochastically and transiently with the MT, dynactin and cargo remained persistently bound and in proximity to the MT. When dynein interacted with a dynactin-cargo complex upon MT-binding, the entire complex moved towards the minus end of MTs in a short run lasting a little over half a second. Repeated rounds of such stochastic interactions of the motor with the MT and cargo complex are required for long-range movement of cargoes. We measured single-molecule parameters pertaining to dynein and used these and other parameters available in literature to develop a MCD model for cargo trajectories based on assumptions from experiments. The experimental data and output from the MCD model for two different kinds of cargo – dextran and EGF vesicles – were in good agreement. Importantly, the experimental data and MCD model output quantitatively agree with the predictions of a 3-state RTP model. Taken together, we demonstrate that stochastic interactions of motors with MTs and cargo complexes are sufficient to elicit complex cargo trafficking behaviour in living cells.

Several interesting points arise from our work. First, the short residence time of dynein that we observed is in contrast with the findings from previous research (McKenney *et al*, 2014; Schlager *et al*, 2014) which report a run length of up to 8 *μ*m for dynein. However, other *in vitro* results have reported dynein run lengths that are comparable with our results (King & Schroer, 2000; Ross *et al*, 2006). It is worth noting that we observed minimal cargo diffusion upon MT depolymerisation, indicative of the highly dense intracellular environment. In *in vitro* assays, the buffer density, as well as factors such as the molar excess of adaptors used, the ionic strength of the buffers, and the source and modifications on the MTs are all likely to differ from the *in cellulo* environment, likely explaining the disparity in dynein run length between our experiments and *in vitro*. Importantly, the run lengths and velocity of cargoes we measure *in cellulo* here are consistent with the well-established trafficking times of endocytic cargo delivery to lysosomes from multiple studies (Rink *et al*, 2005; Salova *et al*, 2017; Kornilova *et al*, 1996; Lakadamyali *et al*, 2006). The short, stop-and-go movements we observe are therefore sufficient for endosomal movement in the cell.

Second, from our study it appears dynactin and endosomal cargoes remain either persistently bound to or remain in very close proximity to MTs. The persistent association of dynactin to MTs that we observed is in contrast to *in vitro* observations where recombinant human dynactin was not found to decorate pig brain-derived MTs (McKenney *et al*, 2014). Similarly recombinant budding yeast dynactin was observed to have a weak interaction with axonemal MTs (Kardon *et al*, 2009). But when dynein was added to the mix, movement of dynactin was observed implying that dynactin could not independently bind to the MTs. However, in other instances (Culver-Hanlon *et al*, 2006), all p150 fragments containing the CAP-Gly domain bound to the MT. Dynactin that was over expressed in cells was found to bind strongly to and bundle MTs (Quintyne *et al*, 1999; Vaughan *et al*, 2002). In fact, dynactin-MT interactions have recently been shown to be important for increasing the on-rate of dynein onto MTs (Sanghavi *et al*, 2021). We propose that dynactin may assist in anchoring cargoes to the MT, allowing rapid initiation of movement upon dynein binding and will investigate this in future studies.

Third, since cytoplasmic dynein is the only minus end-directed motor that participates in membrane trafficking in many cell types, how dynein interacts with, and transports different types of cargo is an interesting question. Recent research suggests that cargo specific adaptors like BicD2 (for Rab6-positive cargo), and Hook1/3 (for Rab5-positive cargo) have differential interaction with dynein (McKenney *et al*, 2014). These cargo adaptors could modulate the interaction time of dynein with the cargo thereby leading to different kinetics. Such stochastic binding and unbinding would allow the same dynein molecule to sample and interact with a wide range of cargo, akin to the ‘Loose Bucket Brigade’ model described previously for kinesin-1 (Blasius *et al*, 2013). Our modelling suggests the differences in cargo movement we observed for dextran and EGF may be due to increased motor on-rates to the cargo. As EGF is highly likely to engage the EGF receptor, while dextran can simply be incorporated into endosomes without receptor binding, we speculate that these differences in motor on-rates are due to the engagement of the EGF receptor, leading to a higher number of motor adaptors being recruited to the endosome. Determining how adaptor number alters endosomal trafficking represents an interesting avenue of future investigation.

Fourth, while we only looked at degradation targeted endosomal cargoes in this study, movement to the MT minus or plus ends (as during cargo recycling) could therefore also be tuned by slightly increasing the bias in one direction, for example by increasing the number of adaptors for dynein on a cargo destined towards the minus end.

Fifth, cellular morphology is likely to play a significant role in the kinetics of motor attachment and detachment from the MT. For instance, in highly polar, narrow axons of neuronal cells, reattachment of dynein to cargo localised close to the MT is likely to occur frequently due to the spatial confinement, as the reduced volume in which dynein could diffuse would lead to higher levels of interaction between the motor and cargo, and therefore more processive cargo trajectories.

Finally, we posit that a similar mechanism is at work in transporting plus end-directed cargo by kinesin motors. More generally, both dyneins and kinesins can simultaneously affect cargo movement, even for those cargo that are transported towards the minus end on the average such as dextran and EGF (Fig. S6i). Further, our modelling suggests the kinesin on-rate for EGF endosomes was also higher than that of dextran endosomes, and the pause times (times that were devoid of both minus and plus end-directed movements) were lower for EGF endosomes compared to dextran. Together, this indicates kinesin binding and subsequent cargo movement may be regulated in a similar fashion to dynein.

Altogether, our study establishes that dynein acts in short spurts in a cellular environment which are sufficient to drive endosomal transport from the cell periphery to perinuclear region within well-established timeframes. Dynein achieves this by stochastic, transient binding to endosomal cargoes, and our modelling suggests cargo adaptor number and/or dynein diffusion upon cargo detachment may be central to determining the ultimate speed of cumulative cargo movement. Future studies experimentally verifying the effects of adaptor number on endosomal movement, refining the role of dynactin in cargo-MT linkages and investigating the behaviour of dynein in polarised cells with constrained cytoplasmic volumes such as neurons and T cells will further shed light on how endosomal movement is controlled.

## Methods

The methods and materials employed in this manuscript are described in detail in the Supplementary Information document.

## ACKNOWLEDGMENTS

We thank LA Chacko for constructive comments on the manuscript, and DH Kim for technical assistance. VA is supported by EMBL Australia and was supported by extramural funding from the Wellcome Trust/Department of Biotechnology–India Alliance (grant IA/18/1/503607), and the Women Excellence Award from the Science and Engineering Re-search Board, India, intramural funding from the Indian Institute of Science, and the RI Mazumdar Young Investigator Award and Department of Science and Technology (India) INSPIRE Faculty Award. KVK acknowledges support from the Department of Atomic Energy, Government of India under project no. RTI4001, the Department of Biotechnology, India, through a Ramalingaswami Re-entry Fellowship and the Max Planck Society through a Max-Planck-Partner-Group at ICTS-TIFR. This research was supported in part by the International Centre for Theoretical Sciences (ICTS-TIFR) for participating in the online program “Thirsting for Theoretical Biology” (code: ICTS/ttb2021/1).

