## Supplementary Information for "Single-molecule imaging of cytoplasmic dynein *in cellulo* reveals the mechanism of motor activation and cargo movement"

**Tirumala, Redpath et al.**

**Vaishnavi**
**Ananthanarayanan.**
****

**This PDF file includes:**

Supporting text
Figs. S1 to S10
Legends for Movies S1 to S10
SI References

**Other supporting materials for this manuscript include the following:**

Movies S1 to S10

### Supporting Information Text

**MCD model.** We consider the stochastic stepping dynamics of two kinds molecular motors (dyneins and kinesins) moving on a MT represented as a one-dimensional lattice of *N* sites with each site being of size $\Delta x$. For simplicity, we assumed periodic boundary conditions on the MT lattice. At each site of the MT, due to the exchange with a large cytoplasmic reservoir, the motors can bind and unbind at rates $[r^D, r^K]$ and $[r^D, r^K]$ respectively. Each site can accommodate a maximum of $M$ motors of any kind. In addition to the motors, there are cargo-units always positioned in close proximity to the MT. There can be only one cargo-unit per lattice site and these units do not have any dynamics of their own. While being bound to the MT at a lattice site with a cargo-unit, the motors can unidirectionally hop along the MT at rates $v_D/\Delta x$ and $v_K/\Delta x$, while dragging the cargo with them. Furthermore, we assumed that (i) all the reactions and lattice-hop processes are Markovian in nature, and that we can model the stochastic dynamics by a master equation for the configuration probabilities, (ii) binding/unbinding events occur at one molecule at a time, (iii) motor(-cargo) hopping is only restricted to the nearest neighbour sites, (iv) the presence of a cargo-complex at a site *does not* affect the kinetics (*i.e.,* the binding/unbinding and movement) of the motors, (v) even in the presence of multiple motors at a site, the motors hop at the same rates, *i.e.,*$v_D/\Delta x$ and $v_K/\Delta x$, and as such, neglect any cooperative binding between multiple motors at a site, (vi) a motor at a site having a cargo-unit at the same site is immediately attached it and can drag along the cargo-complex, (vii) while motors can bind/unbind to the MT independent of the cargo, they can translocate along the MT only if there is a cargo unit at their location on the MT, (viii) the density of cargo-units along the MT track is low and thus we disregarded any steric interactions amongst them, and also that the cargo-units do not diffuse away when not engaged with motors.

At any instant of time *t*, the configuration of this motor-cargo system on the MT is specified by giving the $3N$ integers $(D_1, D_2, \ldots, D_N)$, $(K_1, K_2, \ldots, K_N)$ *and* $(C_1, C_2, \ldots C_N)$ where $D_i, K_i$ *and* $C_i$ are the number of dyneins, kinesins and cargo- units at lattice site $i$. Note that $0 \le D_i \le M$ and $0 \le K_i \le M$ , while each $C_i$ can be either 0 or 1. Periodic boundary conditions imply that $D_{N+1} = D_1$ and $D_0 = D_N$ , and similarly for the $K_i$ and $C_i$. The numbers $D_i$ and $K_i$ at any site $i$ change due to the stochastic binding/unbinding of motors as well as the stochastic hops from neighbouring sites. The cargo numbers $C_i$ change only when motors at *i* hop to the neighbouring sites while dragging the cargo along with them. Note that if $v^D = 0 = v^K$, then each site $i$ would have independent binding/unbinding kinetics. Furthermore, if $M \to \infty$, then the number of dyneins at each site would have an identical steady-state Poisson distribution characterised by a mean number of molecules
$(\langle D \rangle = r^D_{on}/r^D_{off}$ and $\langle K \rangle = r^K_{on}/r^K_{off})$.

Before proceeding further, we define an indicator function

$$\chi_n^M = \begin{cases} 1, & 0 \le n < M \\ 0, & \text{otherwise,} \end{cases} \quad [1]$$

for *n* = 0, 1, 2, 3, . . . and $M \ge 1$. Next, we define $\psi_i$ to be number of dyneins that move from $i+1$ to $i$ as a unit where

$$\psi_i = min(D_{i+1}, M - D_i - K_i), \quad [2]$$

and similarly $\phi_i$ to be number of kinesins that move from $i-1$ to *i* as a unit with

$$\phi_i = min(K_{i-1}, M - D_i - K_i). \quad [3]$$

The joint probability of finding $D_1$ dyneins at lattice-site 1, $D_2$ dyneins at lattice-site 2, . . . , $D_N$ dyneins at lattice-site $N$ , $K_1$ kinesins at lattice-site 1, $K_2$ kinesins at lattice-site 2, . . . , $K_N$ kinesins at lattice-site $N$, $C_1$ cargo-units at lattice-site 1, $C_2$ cargo-units at lattice-site 2, . . . , $C_N$ cargo-units at lattice-site $N$ at time $t$ is denoted by $P\ (D_1, D_2, \ldots, D_N, K_1, K_2, \ldots, K_N, C_1, C_2, \ldots, C_N, t)$.

The master equation governing the temporal evolution of this joint probability is

$$\begin{aligned}
&\frac{d}{dt}P(D_1,D_2,\ldots,D_N,K_1,K_2,\ldots,K_N,C_1,C_2,\ldots,C_N,t)\\
&=\sum_{i=1}^{N}\Bigg[r_{\text{on}}^{D}\,\chi^{M}_{D_i-1+K_i}\,P(\ldots,D_i-1,\ldots,K_i,\ldots,C_i,\ldots)-r_{\text{on}}^{D}\,\chi^{M}_{D_i+K_i}\,P(\ldots,D_i,\ldots,K_i,\ldots,C_i,\ldots)\\
&+r_{\text{off}}^{D}\,(D_i+1)\,P(\ldots,D_i+1,\ldots,K_i,\ldots,C_i,\ldots)-r_{\text{off}}^{D}\,D_i\,P(\ldots,D_i,\ldots,K_i,\ldots,C_i,\ldots)\\
&+r_{\text{on}}^{K}\,\chi^{M}_{D_i+K_i-1}\,P(\ldots,D_i,\ldots,K_i-1,\ldots,C_i,\ldots)-r_{\text{on}}^{K}\,\chi^{M}_{D_i+K_i}\,P(\ldots,D_i,\ldots,K_i,\ldots,C_i,\ldots)\\
&+r_{\text{off}}^{K}\,(K_i+1)\,P(\ldots,D_i,\ldots,K_i+1,\ldots,C_i,\ldots)-r_{\text{off}}^{K}\,K_i\,P(\ldots,D_i,\ldots,K_i,\ldots,C_i,\ldots)\\
&+\frac{v^D}{\Delta x}\,\chi^{M}_{D_i+K_i-\psi_i}\binom{D_{i+1}}{\psi_i}\chi^{1}_{C_i}\,C_{i+1}\,P(\ldots,D_i-\psi_i,D_{i+1}+\psi_i,\ldots,K_i,\ldots,C_i,\ldots)\\
&-\frac{v^D}{\Delta x}\,\chi^{M}_{D_{i-1}+K_{i-1}-\psi_{i-1}}\binom{D_i}{\psi_{i-1}}\chi^{1}_{C_{i-1}}\,C_i\,P(\ldots,D_i,\ldots,K_i,\ldots,C_i,\ldots)\\
&+\frac{v^K}{\Delta x}\,\chi^{M}_{D_i+K_i-\phi_i}\binom{K_{i-1}}{\phi_i}\chi^{1}_{C_i}\,C_{i-1}\,P(\ldots,D_i,\ldots,K_{i-1}+\phi_i,K_i-\phi_i,\ldots,C_i,\ldots)\\
&-\frac{v^K}{\Delta x}\,\chi^{M}_{D_{i+1}+K_{i+1}-\phi_{i+1}}\binom{K_i}{\phi_{i+1}}\chi^{1}_{C_{i+1}}\,C_i\,P(\ldots,D_i,\ldots,K_i,\ldots,C_i,\ldots)\Bigg]
\end{aligned}\qquad \textbf{(S4)}$$

where the first four terms in green represent the binding/unbinding of dynein motors, the next four terms in magenta represent the binding/unbinding of kinesin motors, the next two terms in green represent the translocation of motor-cargo-units due to dynein hops while the last two terms in magenta represent the translocation of motor-cargo-units due to kinesin hops. The binomial coefficients, for instance $\binom{Di+1}{\psi i}$, account for the number of combinations possible for a given hop process. Notice, (i) the motors can bind or hop to a site only if that site can accommodate them, i.e., the maximum number of motors does not exceed $M$ , (ii) the motors can move to the left/right only if they have a cargo-unit at their location and also if the target site does not already have a cargo-unit [these are captured by the factors $\chi^1_{c_i}$ $C_{i+1}$, $\chi^1_{c_i-1}$ $C_i$, $\chi^1_{c_i}$ $C_{i-1}$, and $\chi^1_{c_i+1}$ $C_i$].

As discussed in the main text, we consider the case when a maximum two motors are allowed per site, i.e., $M$ = 2. We simulated the above master using a custom-built python code for the Gillespie algorithm based on the examples in BE150. We explicitly tabulate below the propensity functions for hops of the motor-cargo units shown in Figure S10.

| Hop | Propensity $\times\Delta x$ |
|---|---|
| One dynein hop from site $i$ to $i-1$ | $v^D D_i\,\chi^M_{D_{i-1}+K_{i-1}}\,C_i\,\chi^1_{C_{i-1}}$ |
| Two dyneins hop from site $i$ to $i-1$ | $v^D\,\chi^M_{D_{i-1}+K_{i-1}+1}\,(1-\chi^M_{D_i})\,C_i\,\chi^1_{C_{i-1}}$ |
| One dynein and one kinesin hop from site $i$ to $i-1$ by dynein | $v^D\,\chi^M_{D_{i-1}+K_{i-1}+1}\,D_i K_i\,C_i\,\chi^1_{C_{i-1}}$ |
| One kinesin hop from site $i$ to $i+1$ | $v^K K_i\,\chi^M_{D_{i+1}+K_{i+1}}\,C_i\,\chi^1_{C_{i+1}}$ |
| Two kinesins hop from site $i$ to $i+1$ | $v^K\,\chi^M_{D_{i+1}+K_{i+1}+1}\,(1-\chi^M_{K_i})\,C_i\,\chi^1_{C_{i+1}}$ |
| One dynein and one kinesin hop from site $i$ to $i-1$ by kinesin | $v^K\,\chi^M_{D_{i+1}+K_{i+1}+1}\,D_i K_i\,C_i\,\chi^1_{C_{i+1}}$ |

### Supplementary Methods

**Cell culture.** HeLa cells were cultured in Dulbecco's Modified Eagle Media (DMEM) containing 100 units/mL Penicillin, 0.1 mg/mL Streptomycin, 2 mM L-Glutamine, 400 $\mu$g/mL Geneticin (HiMedia Laboratories) and supplemented with 10% fetal bovine serum (Sigma Aldrich). Cells were grown in an incubator at 37°C under 5% $CO_2$. During live-cell imaging, cells were maintained in live-cell imaging solution (140 mM NaCl, 2.5 mM KCl, 1.8 mM CaCl2, 1 mM MgCl2, 4 mg/mL D-Glucose, 20 mM HEPES) at 37°C using a stage-top incubator (Oko Labs).

**RNAi experiments.** For all RNAi experiments, cells were first grown in glass-bottom imaging dishes (Cat. no. 81518, Ibidi) for 48 h, transfected with the appropriate concentration of siRNA using Jet Prime transfection reagent (Cat. no. 114, Polyplus) and imaged after 48 h. For hDHC RNAi the following siRNA sequence was used: 5'-ACA-UCA-ACA-UAG-ACA-UUC-A-3'10. For p150 RNAi the following siRNA sequences were used: 5'-GUA-CUU-CAC-UUG-UGA-UGA-A-3', 5'-GAU-CGA-GAG- ACA-GUU-AUU-A-3' (1). siRNA oligonucleotides were procured from Eurogentec, Belgium. To quantify the knockdown of proteins, the cells were lysed 48 hours after siRNA transfection to obtain protein solution and SDS-PAGE, western blotting was performed. The following primary antibodies were used to visualise proteins of interest and controls: Rabbit DYNC1H1 Polyclonal Antibody (PA5-49451, Invitrogen, 0.5 $\mu$g/mL), Rabbit Dynactin 1 Polyclonal Antibody (PA5-37360, Invitrogen, 0.5 $\mu$g/mL), Mouse GAPDH Loading Control Antibody (MA5-15738, Invitrogen, 0.5 $\mu$g/mL), Rabbit Actin Polyclonal Antibody (ab8227, AbCam, 0.5 $\mu$g/mL). The following secondary antibodies were used: Anti-Rabbit HRP (31460, Invitrogen, 0.16 $\mu$g/mL) and Anti-Mouse HRP (62-6520, Invitrogen, 0.3 $\mu$g/mL).

**Immunofluorescence.** To immunostain p150 and tubulin, cells were first grown for 48 h in glass-bottom imaging dishes (Cat. no. 81518, Ibidi) and fixed in ice-cold methanol at -20°C for 3 min. Cells were washed thrice in phosphate buffered saline (PBS) for 5 min each and incubated in blocking buffer (5% BSA in PBS) for 60 min. Subsequently, the cells were incubated with p150 and $\alpha$-tubulin antibodies in antibody dilution buffer (1% BSA in PBS) for 60 min. Cells were then washed thrice in PBS for 5 mins and incubated with secondary antibodies in antibody dilution buffer for 45 min. At the end of incubation with secondary antibodies, the cells were washed with PBS and imaged immediately. The following primary antibodies were used: Rabbit Dynactin 1 Polyclonal Antibody (PA5-21289, Invitrogen, 2 $\mu$g/mL), Mouse $\alpha$ Tubulin Monoclonal Antibody (32-2500, Invitrogen, 2 $\mu$g/mL), Rabbit anti-beta Tubulin antibody directly conjugated to AlexaFluor405 (Cat no. EPR16774, Abcam, 250$\mu$g/mL). The secondary antibodies used were Donkey Anti-Rabbit A555 Antibody (A31572, Invitrogen, 0.4 $\mu$g/mL) and Goat Anti-Mouse A647 Antibody (A28181, Invitrogen. 2 $\mu$g/mL). Where anti-beta tubulin was used in conjunction with another rabbit-derived antibody, the other antibody was first added for 2 h at room temperature, detected with an anti-rabbit secondary antibody, sample fixed in 2% PFA for 20 min at room temperature, then anti-beta tubulin AlexaFluor405 added for 1.5 h at room temperature.

To immunostain p150 and p62, p62 and $\alpha$-Tubulin and EB1 and $\alpha$-Tubulin the following method was used. Cells were first grown for 48 h in glass bottomed imaging dishes and fixed in ice-cold methanol at -20°C for 3 min. Subsequently, the cells were washed thrice in PBS for 5 min each and incubated in antibody dilution buffer (2% BSA, 0.1% Triton-X100, in PBS) for 10 min. Next, the cells were incubated in antibody dilution buffer containing the appropriate antibodies for 60 min. Cells were washed thrice in PBS for 5 min each and incubated with appropriate secondary antibodies in antibody dilution buffer for 30 min. At the end of incubation with secondary antibodies, the cells were washed well in PBS and imaged immediately. The following primary antibodies were used. Rabbit Dynactin 1 Polyclonal Antibody (PA5-21289, Invitrogen, 2.5 $\mu$g/mL), Rabbit Alpha Tubulin Monoclonal Antibody (PA5-22060, Invitrogen, 2 $\mu$g/mL), Mouse Dynactin 4 Monoclonal Antibody (MA5-17065, Invitrogen, 2 $\mu$g/mL) and Mouse EB1 Monoclonal Antibody (412100, Invitrogen, 2 $\mu$g/mL). The secondary antibodies used were Donkey Anti-Rabbit A647 Antibody (A32795, Invitrogen, 2 $\mu$g/mL) and Goat Anti-Mouse A555 Antibody (A21422, Invitrogen. 2 $\mu$g/mL).

To immunostain Rab5 and p62, cells were first fixed in 4% Paraformaldehyde (PFA) for 15 mins. The cells were washed well in PBS and incubated for 45 min in antibody staining solution (0.2% Saponin, 0.1% BSA and 0.02% Sodium Azide in PBS) containing primary antibodies against Rab5 and p62. Subsequently, cells were washed well in PBS and incubated for 45 min in antibody staining solution containing appropriate secondary antibodies. Cells were washed well in PBS and imaged immediately. The following primary antibodies were used. Rabbit Rab5 Monoclonal Antibody (3547, Cell Signaling Technology, 14 $\mu$g/mL), Mouse Dynactin 4 Monoclonal Antibody (MA5-17065, Invitrogen, 2 $\mu$g/mL). The secondary antibodies used were Donkey Anti-Rabbit A647 Antibody (A32795, Invitrogen, 2 $\mu$g/mL) and Goat Anti-Mouse A555 Antibody (A21422, Invitrogen. 2 $\mu$g/mL).

**Dextran and EGF uptake.** To visualise fluorescent dextran vesicles, HeLa cells that were grown in glass-bottom imaging dishes (Cat. no. 81518, Ibidi) for 48 h were transferred to serum-free DMEM for 4 h in a 37°C $CO_2$ incubator. Subsequently, the cells were pulsed for 10 min in complete DMEM containing 200 $\mu$g/mL A647-Dextran (D-22914, Invitrogen). Cells were then washed well with live-cell imaging solution before proceeding for microscopy. In all experiments, imaging was completed within 45 min of dextran uptake.

To visualise fluorescent EGF vesicles, HeLa cells that were grown in glass-bottom imaging dishes (Cat. no. P35G-1.5-14-C, Mattek) for 24 h were transferred to serum-free DMEM for 1 h in a 37°C $CO_2$ incubator. Cells were washed in serum-free DMEM, then incubated with 1 nM EGF (Cat. No. E9644, Sigma-Aldrich) labelled with Alexa647 (Cat. No. ab269823, Abcam) in phenol-free, serum-free DMEM for a minimum of 8 minutes prior to imaging. In all experiments, cells were imaged between 8-20 minutes of EGF uptake.

**Transfection.** To visualise fluorescently tagged proteins, cells were transfected with the appropriate plasmids using Jet Prime Transfection Reagent (Cat. no. 114, Polyplus). 3 h after transfection, the cells were washed well in PBS, grown in complete DMEM at 37°C and imaged ~20 h later. The following plasmids were used in this study. (i) mCherry-tubulin was a gift from Mariola Chacon, TUD, Dresden. (ii) Gal4T-mCherry was a gift from Thomas Pucadyil, IISER Pune. (iii) mCherry-DCTN1 was a gift from Kozo Tanaka, Tohoku university, Japan. (iv) mCherry-Rab5 was a gift from Gia Voeltz (Addgene plasmid - 49201). (v) KIF16B-mCherry was a gift from Senthil Arumugam, Monash University, Australia, (vi) Rab7-GFP and LAMP1-GFP were gifts from Jeremie Rossy, University of Konstanz.

**Microscopy.** Imaging was performed on a Nikon Ti2E inverted microscope equipped with a Toptica MLE laser combiner, Nikon H-TIRF Module, Yokogawa CSU-X1 spinning disk module, an Andor iXon 897 EMCCD camera and an Oko Lab stage top incubator. The microscope was controlled using Nikon NIS Elements software or Micromanager (2). Alternately, a Zeiss ELYRA controlled using Zen Black software or a Zeiss Airyscan microscope controlled using Zen Blue software was used.

**HILO microscopy and particle tracking.** For HILO microscopy, a Nikon 100X 1.49NA TIRF objective was used. We optimised the HILO microscopy setting for each cell: First, to avoid overexpression artefacts, we visualised cells expressing low levels of mDHC-GFP. The diameter of the illuminated area was kept constant at 30 $\mu$m and the long axis of cell was aligned perpendicular to the excitation laser. Finally, we adjusted the angle of incidence of the excitation laser, such that the fluorescent spots in a plane ~0.5 $\mu$m from the cover slip appeared bright and distinct. The incidence angle and orientation of excitation laser beam were adjusted using the H-TIRF module and the laser power (50 mW at fiber) was kept at 40%. To visualise single molecules of dynein in Figs. 1, S1d,e, S2 and S4, time-lapse images were acquired at 50 fps with 20 ms exposure per frame for a total of 10 s per video.

To improve the signal-to-noise ratio, a five-frame sliding average of the time-lapse images was used for particle tracking. In these videos, the appearance and subsequent disappearance of intensity at a particular location was classified as a binding event. Such binding events were visually identified and tracked from the start to end frame using Low Light Tracking tool (3) in Fiji/ImageJ (4). The fluorescent spots were classified as stationary, minus-end moving or plus-end moving by marking a region close to the nucleus as the minus end and calculating the displacement of fluorescent spots with respect to the minus end. Particles moving away from the nucleus for >0.32 $\mu$m were classified as plus-end moving, and particles with displacement >0.32 $\mu$m towards the nucleus were minus-end directed, and particles with net displacement <0.32 $\mu$m were classified as stationary. These displacements refer to the total displacement of the dynein particle for the entire duration for which it was tracked. The 0.32 $\mu$m threshold was arrived at after multiplying the 80 nm error in particle tracking with four frames, ensuring that any displacement that is an artefact of the tracking error is discounted in our analysis.

For dual channel imaging of dynein and cargo, videos were acquired at 20 fps with 25 ms exposure per channel for dynein and dextran (in cells depleted of hDHC using RNAi), and at 6 fps with 33 ms exposure per channel for dynein and EGF. To observe dynein-dextran interaction with a better signal-to-noise ratio, a two-frame sliding average of time-lapse images was used.

For 100 s dextran and EGF imaging experiments, an alpha Plan-Apochromat 100x/1.46 Oil Elyra TIRF objective was used. Consistent HILO setting were used across all long-term dextran and EGF uptake experiments. Fluorophores were excited using a 642 nm laser (150mW at source) at a TIRF angle of 63°. Exposure times of 50 ms with a frame interval of 100 ms were used (total frame time, 125 ms with frame transfer), and imaging undertaken for 1000 frames (125 s in total). The movement of dextran vesicles was classified in the following way: the movement between two consecutive frames was classified as zero if it was less than 20 nm (the tracking accuracy of low light tracking tool under the imaging conditions used (3).). Then, displacement for four consecutive frames towards or away from the nucleus was classified as a minus or plus end-directed run, and a zero displacement for 2 or more consecutive frames was classified as a pause. The same criteria for runs and pauses was used for analysing the trajectories obtained from the MCD model.

**Spinning disk (SD) confocal microscopy and image analysis.** To quantify the correlation between expression level of dynein and its clustering, z-stack images of live HeLa cells expressing mDHC-GFP were acquired using a 60x, 1.4 NA objective with 50 ms exposure. The intensity comparison was done for the lowest plane at which clusters were distinctly visible. The dynamics of mCherry-p150 in cells were visualised by acquiring 60-s long movies of the cell with a 1.4 NA 60x objective, 100 ms exposure per frame and 1 s interval between frames. The dynamics of dextran vesicles along the MTs was visualised by acquiring 30-s long movies of the cell with a 60x, 1.4 NA objective, 100 ms sequential exposure/channel and 1 s interval between frames.

The SD+SRRF images were obtained by taking the mean of the radiality map of 100 images of a single field acquired under a 100x, 1.49NA objective with the SD confocal microscopy setup. The radiality magnification in SRRF was set to 4 for all experiments. To visualise the association between MTs and dextran vesicles in live cells, the SRRF acquisition settings used were 20ms exposure, ring radius of 0.5 for MTs and 3 for dextran. To visualise p62 with tubulin and EB1 with tubulin, 20 ms exposure and ring radius of 0.5 was used. To visualise p150 with tubulin, and p150 with p62 the SRRF acquisition settings used were 100 ms exposure, ring radius of 1. Finally, to visualise p62 with Rab5, 20ms exposure was used in both channels. A ring radius of 0.5 and 3 were used for p62 and Rab5, respectively.

Kymographs for all live-cell images were generated as follows: first, the cell was oriented so that its long axis was vertical (horizontal) within the x-y plane of the image window. The 'Reslice' function in Fiji/ImageJ was applied starting left (top), rotate 90° (option not selected). Then a maximum intensity projection was obtained to selectively visualise trajectories along the long axis.

The cooccurrence of signals with each other (e.g. p150 with p62) was quantified by first thresholding SRRF images of p150 for the top 5% of intensity. The number of p150 spots was counted $N_{p150}$, and a mask was created. Then, the SRRF image of p62 in the same cells was combined with the p150 mask using AND operation in Fiji. The resultant image was thresholded and the number of spots counted ($N_{p62}$). $N_{p62}/N_{p150}$*100 gave the percentage of p150 spots with p62. A similar procedure was used to quantify the percentage of cooccureence of all other signals. To quantify the effect of p150 knock-down on the levels of p150, p62 along the MT, ROIs were drawn along distinct MT segments that were randomly chosen in SRRF images of MT, and the mean intensities of p150 or p62 in these ROIs were measured.

**Correlative Light and Electron Microscopy.** HeLa cells were grown for 24 hours on gridded glass-bottom imaging dishes (Cat. No. P35G-1.5-14-CGRD, Mattek), following which cells were transfected with EGFP-tubulin (Cat. No. 56450, AddGene) overnight. Cells were transferred into serum-free, phenol free DMEM for 1 h, following which A647-dextran or EGF-647 were added to cells for 10 min. Cells were extensively washed in PBS and fixed in 4% PFA containing 0.2% glutaraldehyde (Cat. No. 354400, Sigma-Aldrich) for 1 h at room temperature. Following fixation, cells were dyed with MitoTracker Orange CMTMRos (Cat. No. M7510) to fluorescently label mitochondria through the cell volume, then imaged on a Zeiss LSM 800 with Airyscan detector, with a pixel size of 42.5 nm and z-slice spacing of 150 nm. The MitoTracker-stained mitochondria served as markers for correlation and registration of the relevant confocal z-slice with that of the corresponding TEM z-slice based on identification of mitochondrial morphology and distribution at each slice (Fig. S9). After imaging, cells were further fixed in 2.5% glutaraldehyde in 0.1 M sodium cacodylate for 1 h at room temperature. All the subsequent processing steps were carried out in a BioWave microwave (Pelco). Post fixation, samples were washed with cacodylate buffer, additionally fixed in reduced osmium (1% osmium tetroxide-1.5% potassium ferricyanide) solution followed by buffer wash, resuspended in 2% aqueous $OsO_4$ (osmium tetroxide) (ProScitech). Washed samples were then stained with 2% w/v Phosphotungstic acid (PTA) in 30% ethanol at 60° for 1 h, followed with a 30% ethanol wash at room temperature. Samples were then stained with 2% aqueous uranyl acetate prepared in 30% ethanol, incubated at 4°C for 1 h. Serial dehydration was then continued at this point in increasing percentages of ethanol, following which cells were serially infiltrated with Durcupan ACM (Cat. No. 44610, Sigma). Fresh 100% resin was then added and polymerised at 60°C overnight. Ultrathin sections were cut on an ultramicrotome (UC6: Leica) and imaged at 100 kV on a JEOL1400 transmission electron microscope fitted with a Phurona EMSIS CMOS camera using a 5x5 stitching matrix. Light and electron microscopy images were subsequently overlayed and correlated using the Correlia plugin for Fiji/ImageJ (5) and is briefly described here. Using the signal from the orange fluorescent mitochondrial dye, a TEM section of the cell 65-nm thick was correlated with the best fit confocal z-slice within the realms of the confocal slice thickness of 150 nm. An overlay of the TEM image and corresponding confocal image showing the mitochondrial signal only at the relevant z-slice was created for cells that had taken up both A647-dextran and A647-EGF. Once the overlay confirmed the right cell volume, then the green signal from EGFP-tubulin and the red signal from A647-dextran or EGF-647 were overlayed onto the same TEM image and this enabled visualisation of the proximity of dextran and EGF endosomes with respect to MTs.

**Curve fitting and data visualisation.** For the intensity fits in Fig. 1c, the intensity values of tracked dynein molecules was plotted as a histogram. This distribution was fit by a sum of two Gaussians with the same standard deviation ($\sigma$). The smaller mean was constrained between 15-30 and the larger mean was constrained between 35-50.

For the exponential fit in Fig. 1f, the probability vs residence time ($\tau$ ) histogram of dynein on the MTs was obtained from the lengths of all single molecule binding events. From this $P(\tau)$= 1-cumulative frequency of the residence time was obtained, and was fit to $\lambda e^{-\lambda t}$ to obtain $\lambda$, the off-rate from MTs. The average residence time was given by $1/\lambda$. The fits in Figs. 7, 8 and 9 were similarly obtained using in-built Python functions.

All data was plotted using MATLAB (MathWorks) or Python. Statistical analyses were performed in Matlab and the specific tests performed are indicated in the respective figure captions. The statistical tests or standard deviation calculations were performed with the number of data points referring to individual cells or endosomes (n) pooled over multiple independent experiments (N). Figures panels were prepared using Adobe Illustrator.

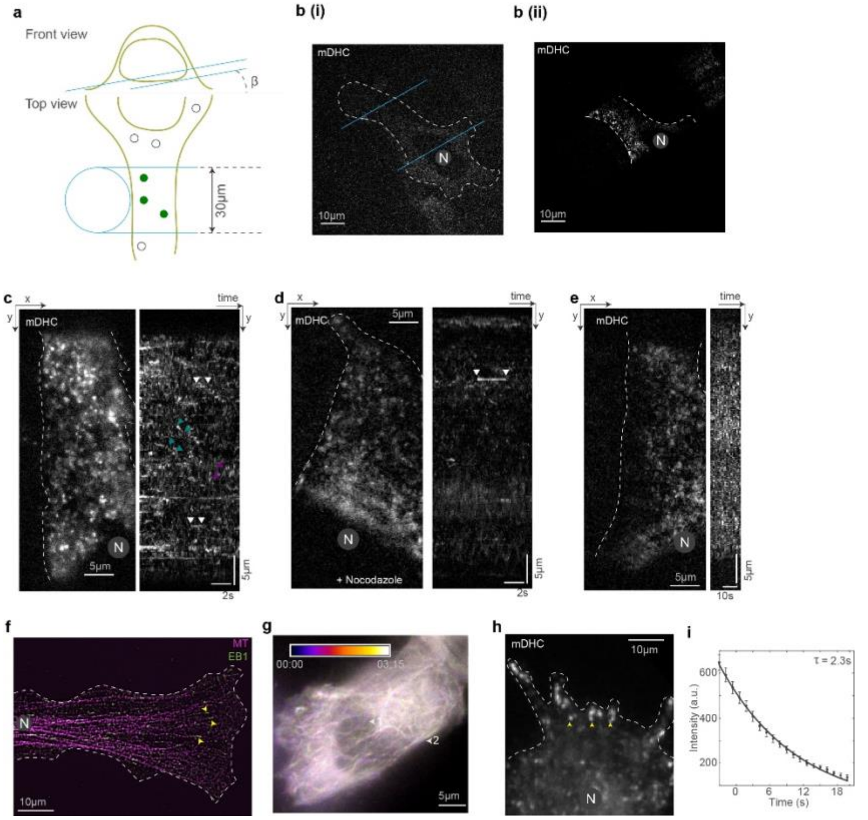


**Fig. S1. Visualisation of single molecules of dynein using HILO microscopy. a**, HILO microscopy setup used to visualise dynein molecules in HeLa cells. Depending on the morphology of the cell, the angle $\beta$ was adjusted such that fluorescent spots of dynein were visible (top, 'Front view'). The illumination diameter was kept constant at 30 $\mu$m using a field stop (bottom, 'Top view'). This resulted in a beam thickness of ~4.6 $\mu$m (calculated according to (6)). **b**, (i) SD confocal microscopy image of a HeLa cell expressing mDHC-GFP and (ii) HILO microscopy image of the same cell, that is partially illuminated in the HILO microscopy setup. The blue lines represent the orientation of the incident laser beam. In the HILO microscopy image, individual fluorescent spots are visible. **c**, HILO microscopy image (left) from a 10-s long time-lapse video of mDHC-GFP in HeLa cells and the corresponding kymograph (right). Representative stationary, minus end-directed and plus end-directed events are indicated with the white, teal and magenta arrowheads respectively in the kymograph. **d**, HILO microscopy image (left) from a 10-s long time lapse of mDHC-GFP in cells treated with 10 $\mu$M nocodazole to depolymerise the MTs and the corresponding kymograph (right). The kymograph shows few binding events compared to the cell shown in **c** indicating that dynein molecules were stochastically binding to the MTs from the cytosol. **e**, HILO microscopy image (left) from a 20-s long 1 fps time lapse video of mDHC-GFP in HeLa cells and the corresponding kymograph (right). There are no distinct traces in the kymograph indicating that the short traces shown in **d** are not artefacts and dynein molecules do not interact with the MTs for a long duration. **f**, Immunofluorescence images of MT (magenta) and EB1 (green) obtained using SD microscopy + SRRF. In these cells with a large aspect ratio, a majority of the MTs are plus end out (pointed by yellow arrow heads). In all analysis, movement towards the nucleus was considered as minus end-directed transport and movement away from the nucleus as plus end-directed. **g**, Temporally colour-coded signal from mCherry-tubulin in a 3min 15s video obtained using HILO microscopy. This indicated that the MTs position and dynamics did not change significantly in the course of the movie. Moreover, a significant length (~20 $\mu$m) of the MT network was visible in HILO microscopy images indicating that dynein molecules moving on MTs over long distances could be visualised and tracked. **h**, HILO microscopy image from a time-lapse video of mDHC-GFP in a cell expressing high levels of mDHC-GFP. Clusters of dynein that are likely at the MT plus end are indicated by the yellow arrowheads. **i**, An exponential decay fit to the intensity vs time plot of spots similar to those indicated with yellow arrowheads in g indicating that a dynein molecule with intensity ~23 a.u. could be visualised for ~50 s before it was photobleached (23 x 2.3). Data from n =119 spots, N=1 independent experiment with 25 cells. In **b**-**e**, **f** and **h**, 'N' marks the location/direction of the nucleus.

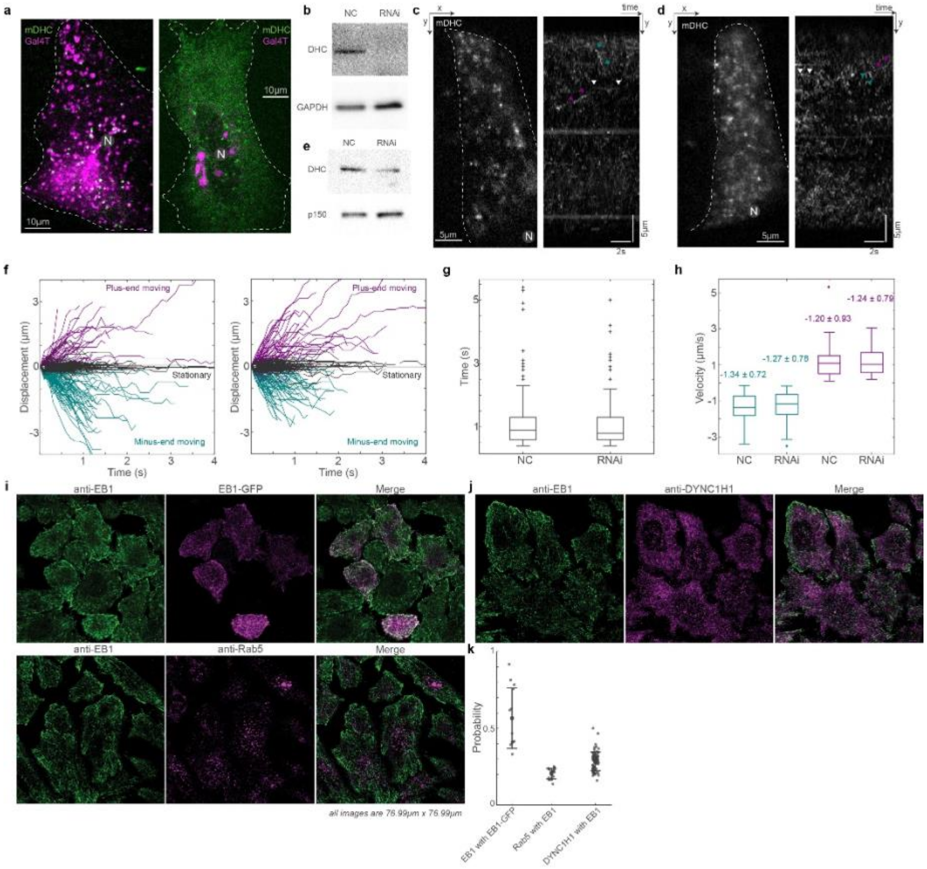


**Fig. S2. mDHC-GFP is functional in HeLa cells depleted of endogenous DHC. a**, SD microscopy images of live cells expressing Gal4T-mCherry (magenta) and mDHC-GFP (green). The cells were treated with 35 nM siRNA against endogenous hDHC. The cell on the left had no discernible mDHC-GFP signal and a dispersed Golgi apparatus whereas the cell on the right expressed mDHC-GFP and had the Golgi clustered at the cell center. Quantification revealed that 100% of the cells without mDHC-GFP expression had a dispersed Golgi whereas only 24% of the cells expressing mDHC-GFP had a dispersed Golgi, indicating mDHC-GFP was functional (n=34 cells from N=2 independent experiments). **b**, Representative western blot to verify the knock down of hDHC by RNAi. Quantification of the western blot confirmed that the RNAi successfully knocked down levels of hDHC by >90% (2 independent experiments). **c**, HILO microscopy image (left) from a 10-s long time lapse of mDHC-GFP in cells treated with 25 nM negative control (NC) siRNA and the corresponding kymograph (right). Representative stationary, minus end-directed and plus end-directed events are indicated with white, teal and magenta arrowheads respectively in the kymograph. **d**, HILO microscopy image (left) from a 10-s long time-lapse of mDHC-GFP in cells treated with 25 nM hDHC siRNA and the corresponding kymograph (right). Representative stationary, minus end-directed and plus end-directed events are indicated with white, teal and magenta arrowheads respectively in the kymograph. **e**, Representative western blot to verify the knockdown of DHC by RNAi. Quantification of the western blot confirmed that the RNAi successfully knocked down levels of DHC by an average of 72% (2 independent experiments). **f**, Comparison of displacement vs time plots of mDHC-GFP molecules in cells treated with 25 nM NC siRNA (left) and 25 nM siRNA against endogenous DHC (right). In cells treated with 25 nM NC siRNA, 53% of the molecules remained stationary, 31% moved towards the minus ends of the MTs and 16% moved towards the plus ends of the MTs, whereas in cells treated with 25nM siRNA against hDHC, 52% of the molecules remained stationary, 29% moved towards the minus ends of the MTs and 19% moved towards the plus ends of the MTs. **g**, Comparison of the residence time of mDHC-GFP molecules in cells treated with 25 nM NC siRNA vs cells treated with 25 nM siRNA against endogenous hDHC indicating no significant differences. **h**,Comparison of minus end-directed velocities (teal boxes) of mDHC-GFP molecules in cells treated with 25 nM NC siRNA and 25 nM siRNA against endogenous DHC, showing no significant differences. Similarly, the plus end-directed velocities (magenta boxes) were also not significantly different. The mean ±s.d. of velocities is indicated in the box plot. In **f**-**h** data for NC was obtained from n=254 binding events tracked from N=3 independent experiments with >30 cells. Data for RNAi was obtained from n=245 binding events tracked from N=3 independent experiments with >30 cells. **i**, Fluorescence images of anti-EB1 (left, green), EB1-GFP (magenta, top centre), anti-Rab5 (magenta, bottom centre), and their merge (right), obtained using Airyscan confocal microscopy. **j**, Top - fluorescence images of anti-EB1 (left, green), anti-DYNC1H1 (magenta, centre), and their merge (right) obtained using Airyscan confocal microscopy. **k**, Quantification (mean ±s.d.) of the co-occurrence of the different proteins imaged in **i** and **j**. Note that 'EB1 with EB1-GFP' is a positive control and represents the maximum co-occurrence that is quantifiable in cells, 'Rab5 with EB1' is a negative control, since EB1 and Rab5 do not typically interact. Each dot represents an individual cell analysed (n=13-74 cells from N>=2 independent experiments). In **a**, **c**, and **d**, 'N' marks the location/direction of the nucleus.

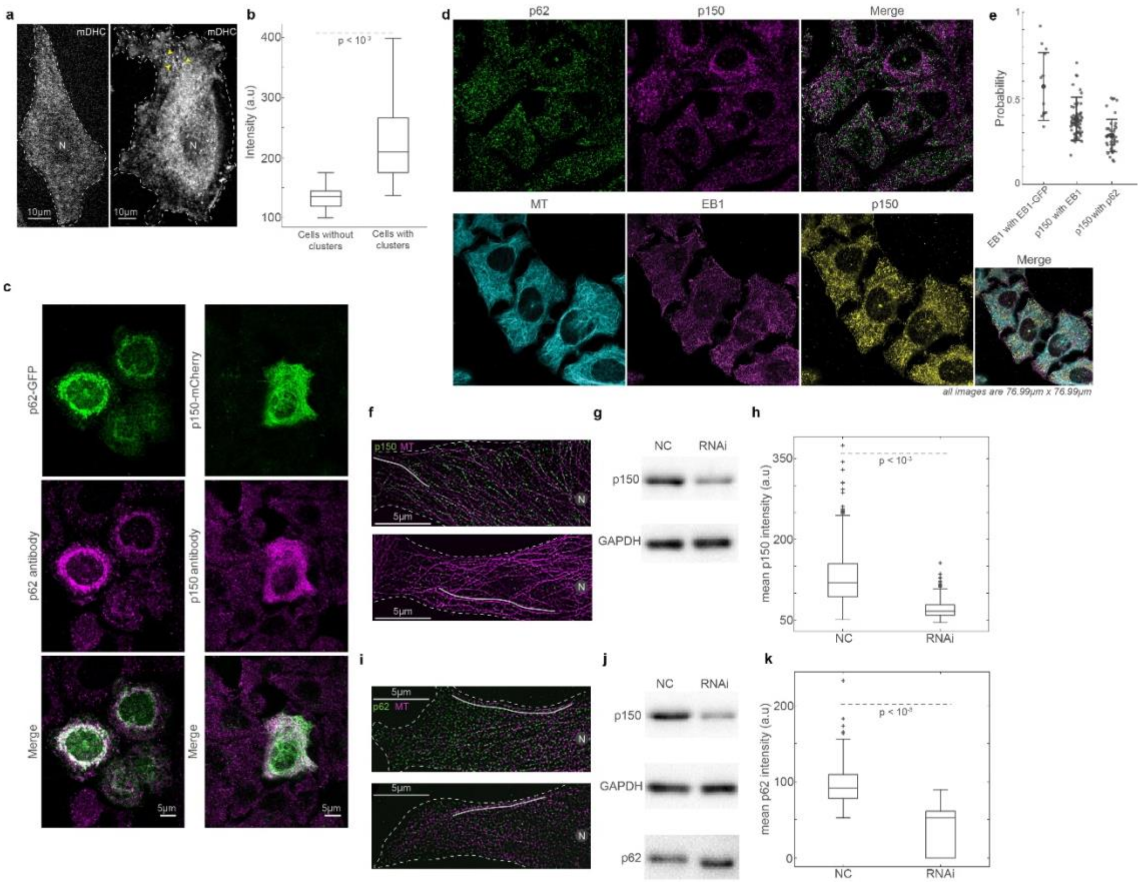


**Fig. S3. Dynactin's association with MTs is long-lived.** **a**, SD microscopy images of live cells expressing mDHC-GFP, with no visible dynein clusters (left) and with distinct dynein clusters (yellow arrow heads) (right). **b**, Comparison of average intensities between cells with and without dynein clusters showing that cells with higher total mDHC intensities exhibited clusters. Therefore, clusters of dynein were likely a result of high over expression of mDHC. (n=25 cells per condition from N=1 independent experiment). **c**, Left - fluorescence images of p62-GFP (top, green), anti-p62 (magenta, middle), and their merge (bottom). Right - fluorescence images of p150-GFP (green, top), anti-p150 (Invitrogen antibody, magenta, middle), and their merge (bottom). These images indicate that the antibodies employed to detect p62 and p150 in this study are specific. **d,** Top - Immunofluorescence images of p62 (left, green), p150 (detected using the Invitrogen antibody, 'p150[1]', centre, magenta), and their merge (right). Bottom - Immunofluorescence images of MT (left, cyan), EB1 (second from left, magenta), p150 third from left, yellow) and their merge (right). All images were obtained using Airyscan confocal microscopy. **e**, Quantification (mean $\pm$s.d.) of the co-occurrence of the different proteins imaged in **d**. Note that 'EB1 with EB1-GFP' has been reused from Fig. S2k. Each dot represents an individual cell analysed (n=45-74 cells from N>=2 independent experiments). All images were obtained using Airyscan confocal microscopy. **f**, Immunofluorescence images of MT (magenta) and p150 (green) obtained using SD microscopy + SRRF on cells treated with 35 nM NC siRNA (top) and 35 nM p150 siRNA (bottom). **g**, Representative western blot to verify the knock down of p150 by RNAi. Quantification of the western blot confirmed that the RNAi successfully knocked down levels of p150 by an average of 60% (2 independent experiments). **h**, Comparison of mean intensity of p150 along MTs (representative ROI shown as grey line in **f** between cells treated with NC and p150 siRNA showing that p150 knockdown reduces p150 levels along MTs. For each condition, 500 MT segments of length $15 \pm 6$ $\mu$m (mean $\pm$s.d.) were analysed from n=50 cells from N=2 independent experiments. **i**, Immunofluorescence images of MT (magenta) and p62 (green) obtained using SD microscopy + SRRF on cells treated with 35 nM NC siRNA (top) and 35 nM p150 siRNA (bottom). **j**, Representative western blot to verify the knock down of p150 by RNAi. Quantification of the western blot confirmed that the RNAi successfully knocked down levels of p150 by an average of 60% (2 independent experiments). Moreover, there were no significant differences in the levels of p62 in cells treated with NC and p150 siRNA. **k**, Comparison of mean intensity of p62 along MTs (representative ROI shown as grey line in **i** between cells treated with NC and p150 siRNA showing that p150 knockdown reduces p62 levels along MTs. For NC, ~190 MT segments of length $15 \pm 6$ $\mu$m (mean $\pm$s.d.) were analysed from n=51 cells from N=2 independent experiments. For p150 RNAi, ~100 MT segments of length $17 \pm 9$ $\mu$m (mean $\pm$s.d.) were analysed from n=52 cells from N=2 independent experiments. In **a**, **f**, and **i**, 'N' marks the location/direction of the nucleus.

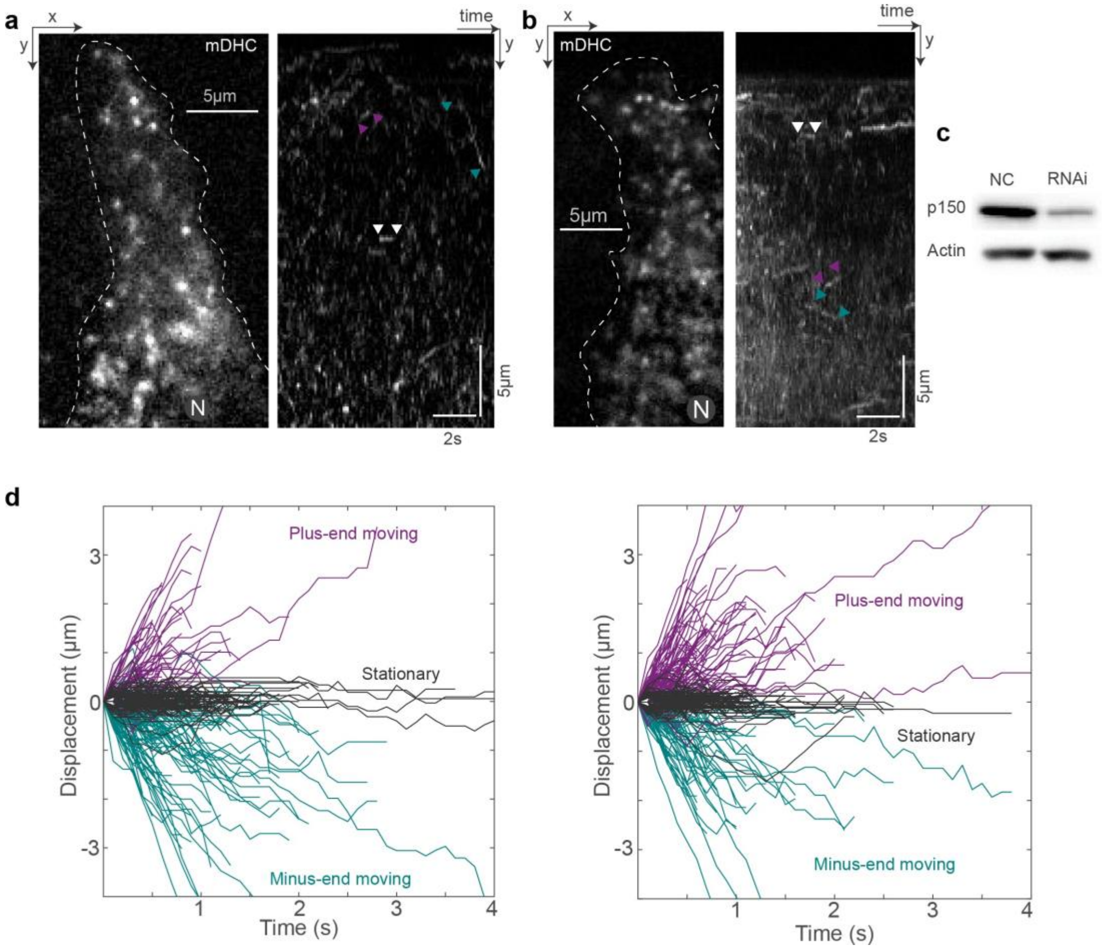


**Fig. S4. The effect of dynactin depletion on dynein behavior.** **a**, HILO microscopy image (left) from a 10-s long time lapse video of mDHC-GFP in cells treated with 25 nM NC siRNA (left) and the corresponding kymograph (right). Representative stationary, minus end-directed and plus end-directed events are indicated with white, teal and magenta arrowheads respectively in the kymograph. **b**, HILO microscopy image (left) from a 10-s long time lapse video of mDHC-GFP in cells treated with 25 nM siRNA against endogenous p150 and the corresponding kymograph (right). Representative stationary, minus end-directed and plus end-directed events are indicated with white, teal and magenta arrowheads respectively in the kymograph. **c**, Representative western blot to verify the knockdown of p150 by RNAi. Quantification of western blot confirmed that the RNAi successfully knocked down levels of p150 by an average of 68% (2 independent experiments). **d**, Comparison of displacement vs time plots of single dynein molecules in cells treated with 25 nM NC siRNA (left) and 25 nM siRNA against p150 (right). In cells treated with 25 nM NC siRNA, 48% of the molecules remained stationary, 34% moved towards the minus ends of the MTs and 18% moved towards the plus ends of the MTs, whereas in cells treated with 25 nM siRNA against p150, 50% of the molecules remained stationary, 21% moved towards the minus ends of the MTs and 29% moved towards the plus ends of the MTs. Data for NC obtained from n=274 binding events tracked from N=2 independent experiments with >30 cells each. Data for RNAi obtained from n=299 binding events tracked from N=2 independent experiments with >30 cells each. In **a** and **b**, 'N' marks the location/direction of the nucleus.

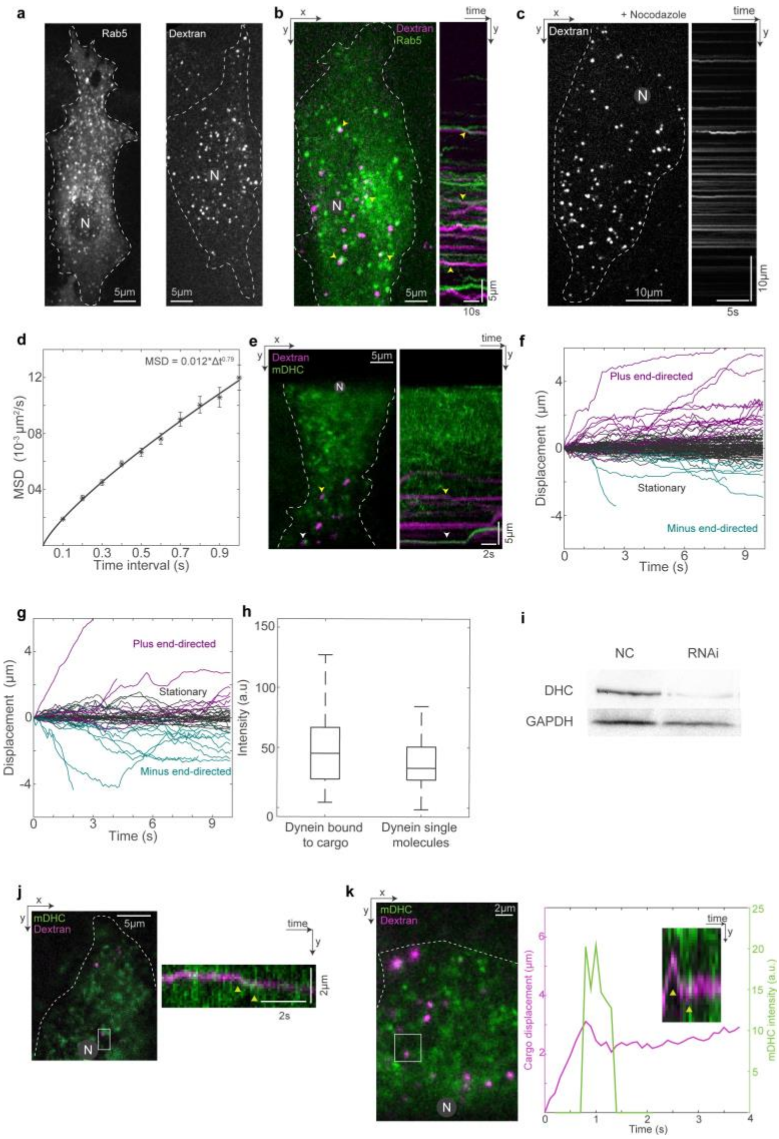


**Fig. S5. Dextran as endosomal cargo marker and its interaction with dynein. a**, SD microscopy images of mCherry-Rab5 (left) and dextran-A647 (right). The signal-to-noise ratio was higher in cells with endocytosed dextran allowing us to track dextran vesicles with high spatio-temporal resolution and perform dual channel imaging along with single molecules of dynein. **b**, SD microscopy image from a 60-s long time lapse video of mCherry-Rab5 (green) and dextran-A647 (magenta) in cells (left) and the corresponding kymograph (right). Yellow arrow heads point to colocalised Rab5 and dextran indicating dextran vesicles were a proxy for early endosomal compartments. In images acquired within 60 min after a 10-min pulse of dextran, 63 ± 14% of the dextran vesicles were associated with a Rab5 punctae (n=125 dextran vesicles from N=1 independent experiment with 17 cells). **c**, SD microscopy image from a 10-s long time lapse video of dextran-A647 in cells treated with 10 $\mu$M nocodazole (left) and the corresponding kymograph (right). The kymograph shows abrogation of directed transport, as expected upon MT depolymerisation. **d**, Mean squared displacement (MSD) analysis of dextran vesicles tracked in cells treated with 10 $\mu$M nocodazole for >30 mins. The mean squared displacement data of dextran vesicles was fit to $\alpha *(\Delta t)\beta$ and the coefficient $\alpha$ was estimated to be 0.012 $\mu m^2$/s indicating that even in the absence of MTs, intracellular crowding likely prevented the dextran vesicles from diffusing away. (n=804 dextran vesicles from N=1 independent experiment with 24 cells). **e**, HILO microscopy image (left) from a 10-s long time lapse video of mDHC-GFP (green) and dextran-A647 (magenta) and the corresponding kymograph (right). The yellow arrowhead points to a dextran vesicle with no dynein fluorescence and the white arrowhead points to a dextran vesicle with associated dynein intensity. The kymograph shows that dextran vesicle with dynein bound to it (white arrowhead), made a short run towards the MT minus end. The dextran vesicle without visible dynein intensity (yellow arrowhead) remained stationary. **f**, Displacement vs time plot of dextran vesicles without associated dynein intensity. **g**, Displacement vs time plot of dextran vesicles with visible dynein intensity.Comparing **f** and **g**, vesicles bound to dynein had a higher likelihood (28%) of moving towards the MT minus ends than dextran vesicles not associated with dynein (10%) (n=158 dextran vesicles from N=2 independent experiments with >25 cells). **h**, Comparison of intensity of dynein on dextran vesicles and intensity of single molecules of dynein (measured from binding events) showed that there were 1-2 dynein molecules per dextran vesicle (n=44 dextran vesicles with dynein intensity, n=115 single molecules of dynein from N=2 independent experiments with >25 cells). **i**, Analysis **f**-**h** was done in cells expressing mDHC-GFP and treated with 25 nM siRNA against endogenous hDHC. Quantification of western blot confirmed that the RNAi successfully knocked down levels of DHC by 85%. **j**, HILO microscopy image (left) from a time lapse video of mDHC-GFP (green) and dextran-A647 (magenta), and the kymograph corresponding to the region marked with the white square (right). The yellow arrowheads point to a short run of the dextran vesicle towards the MT minus end upon binding of a dynein molecule. **k**, HILO microscopy image (left) from a time lapse video of mDHC-GFP (green) and dextran-A647 (magenta) in cells with RNAi-mediated depletion of hDHC, and the kymograph and plots corresponding to the region marked with the white square (right). The yellow arrowheads point to a short run of a previously plus end-directed dextran vesicle towards the MT minus end upon binding of a dynein molecule. In **a**, **b**, **c**, **e**, **I, j and k**, 'N' marks the location/direction of the nucleus.

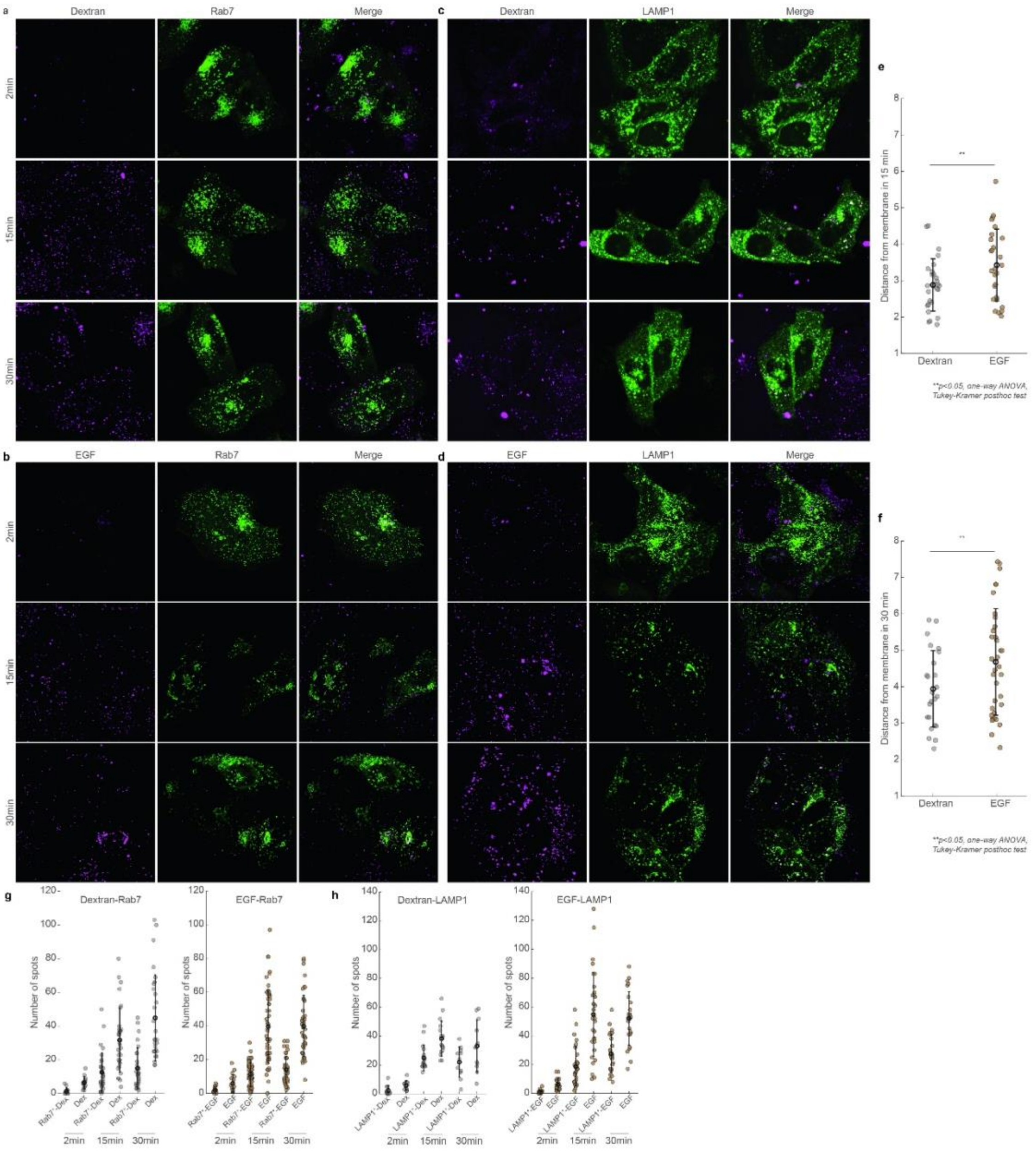


**Fig. S6. Net movement of dextran and EGF endosomes over time.** **a**, Airyscan confocal images of dextran-AF647 (magenta, left), Rab7-GFP (green, centre) and their merge in HeLa cells fixed 2 min (top), 15 min (middle) and 30 min (bottom) after introduction of dextran to the cells. **b**, Airyscan confocal images of EGF-AF647 (magenta, left), Rab7-GFP (green, centre) and their merge in HeLa cells fixed 2 min (top), 15 min (middle) and 30 min (bottom) after introduction of EGF to the cells. **c**, Airyscan confocal images of dextran-AF647 (magenta, left), LAMP1-GFP (green, centre) and their merge in HeLa cells fixed 2 min (top), 15 min (middle) and 30 min (bottom) after introduction of dextran to the cells. **d**, Airyscan confocal images of EGF-AF647 (magenta, left), LAMP1-GFP (green, centre) and their merge in HeLa cells fixed 2 min (top), 15 min (middle) and 30 min (bottom) after introduction of EGF to the cells. **e**, Quantification (mean ± s.d.) of the distance of dextran and EGF vesicles from the membrane in cells fixed 15 min after the introduction of AF647-labelled dextran or EGF. Each dot represents the mean distance of several endosomes from an individual cell (n=29 for dextran data, and n=29 cells for EGF data, from N=3 independent experiments). **f**, Quantification (mean ± s.d.) of the distance of dextran and EGF vesicles from the membrane in cells fixed 30 min after the introduction of AF647-labelled dextran or EGF. Each dot represents the mean distance of several endosomes from an individual cell (n=26 cells for dextran data, and n=37 cells for EGF data, from N=3 independent experiments). **g**, Plot of the number of Rab7$^+$ dextran vesicles compared to the number of dextran vesicles (left), and plot of number of Rab7$^+$ EGF vesicles compared to the number of EGF vesicles (right) at 2 min, 15 min and 30 min after introduction of AF647-labelled dextran or EGF to HeLa cells. **h**, Plot of the number of LAMP1$^+$ dextran vesicles compared to the number of dextran vesicles (left), and plot of number of LAMP1$^+$ EGF vesicles compared to the number of EGF vesicles (right) at 2 min, 15 min and 30 min after introduction of AF647-labelled dextran or EGF to HeLa cells.

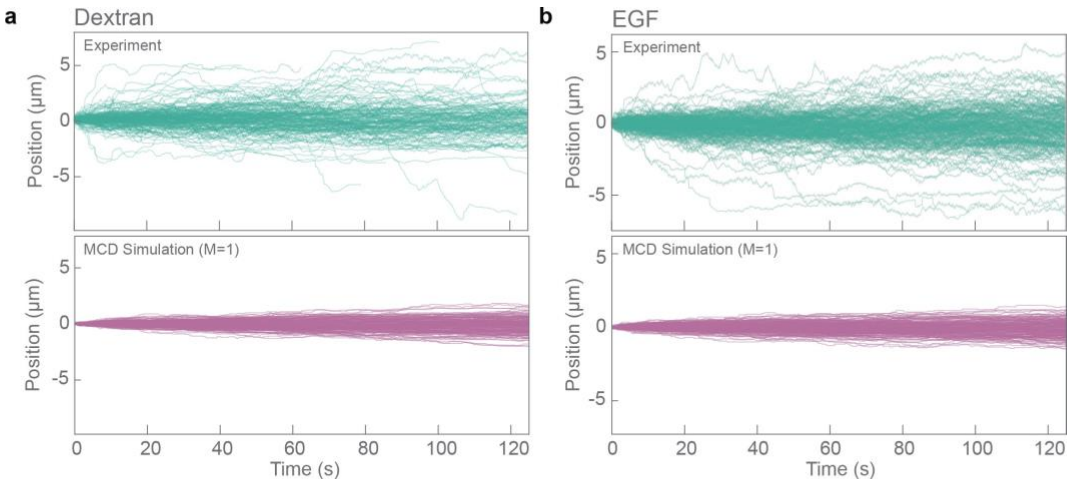


**Fig. S7. Trajectories from the MCD model when $M$ =1 a**, Position vs. time trajectories of dextran vesicles from experimental data (green, top) and from the MCD simulation (magenta, bottom), when $M = 1$, i.e., we did not allow the possibility of pairs of motors to transport cargo. **b**, Position vs. time trajectories of EGF vesicles from experimental data (green, top) and from the MCD simulation (magenta, bottom), when $M = 1$.

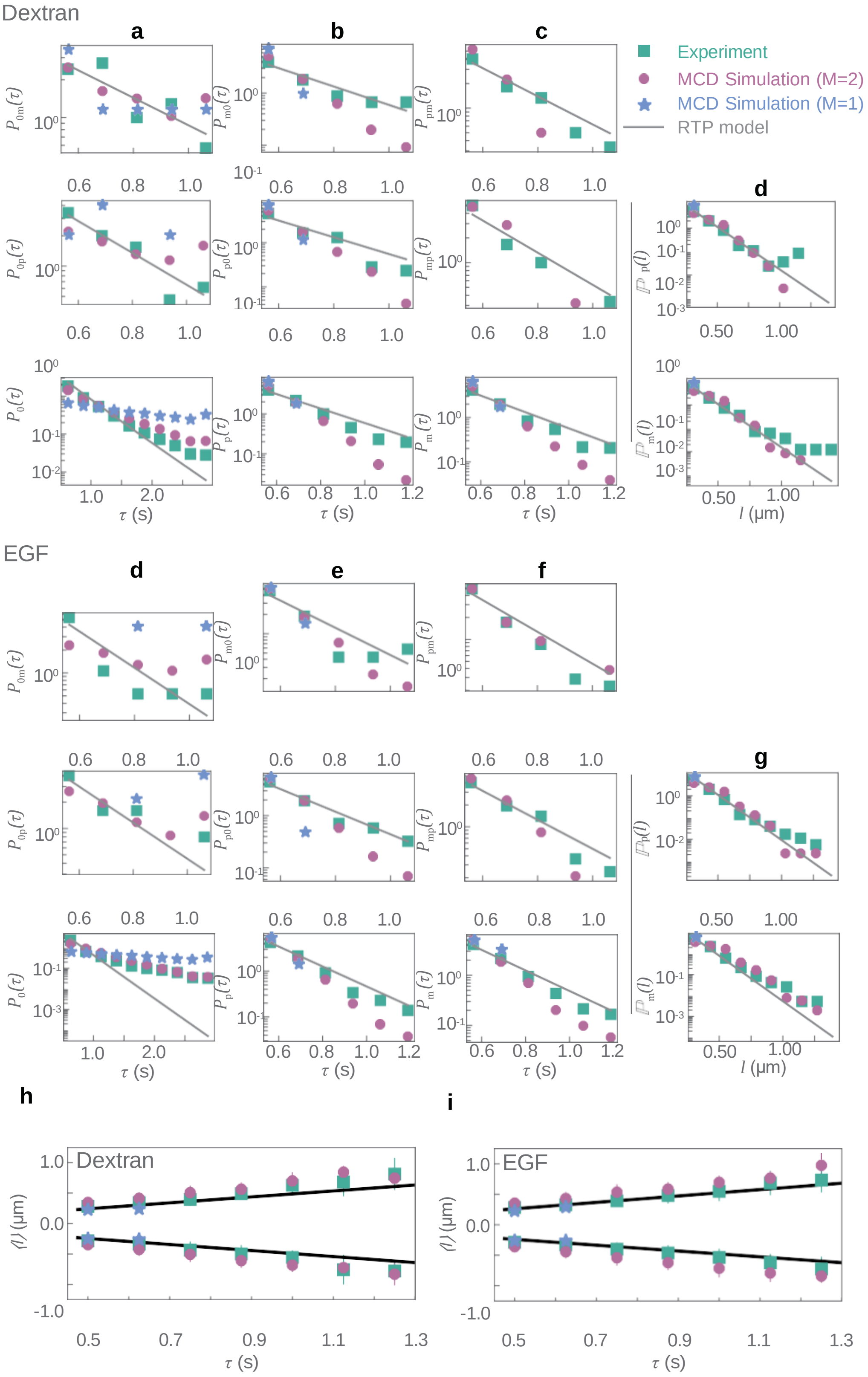


**Fig. S8. RTP fits for dextran and EGF trajectories**, Probabilities distributions of the waiting time for cargo movement from stationary to minus end-directed (**a** top, **d** top), minus end-directed to stationary (**b** top, **e** top), plus to minus end-directed (**c** top, **f** top), stationary to plus end-directed (**a** middle, **d** middle), plus end-directed to stationary (**b** middle, **e** middle) and minus end-directed to plus end-directed (**c** middle, **f** middle) are plotted against the time interval ($\tau$) for data from experiments (green squares) and output from the MCD simulations (magenta circles). The probability of the residence times of cargo in the stationary state (**a** bottom, **d** bottom), in the plus end-directed state (**b**

bottom, **e** bottom), and minus end-directed state (**c** bottom, **f** bottom) are plotted against the time interval ($\tau$) for data from experiments (green square) and output from the MCD model with $M = 2$ (magenta circles) and $M = 1$ (blue stars). Distribution of the run lengths in the plus end-directed (**d** top, **g** top) and minus end-directed states ((**d** bottom, **g** bottom) plotted against the run lengths ($l$) for data from experiments (green square) and output from the MCD model with $M = 2$ (magenta circles) and $M = 1$ (blue stars). The grey lines represent fits to the experimental data from the RTP model. Please note that plots **a**-**g** are in log-linear scale. Plot of average run length ($\langle l \rangle$) as a function of run time ($\tau$) for dextran vesicles (**h**) and EGF vesicles (**i**), with data from experiments (green square) and output from the MCD model with $M = 2$ (magenta circle) and with $M = 1$ (blue stars). The black solid line represents the prediction for experimental data from the RTP model. Note that the experimental data and output from MCD model with with $M = 2$ are identical to those in Fig. 7, 8 and 9, and are reproduced here for comparison.

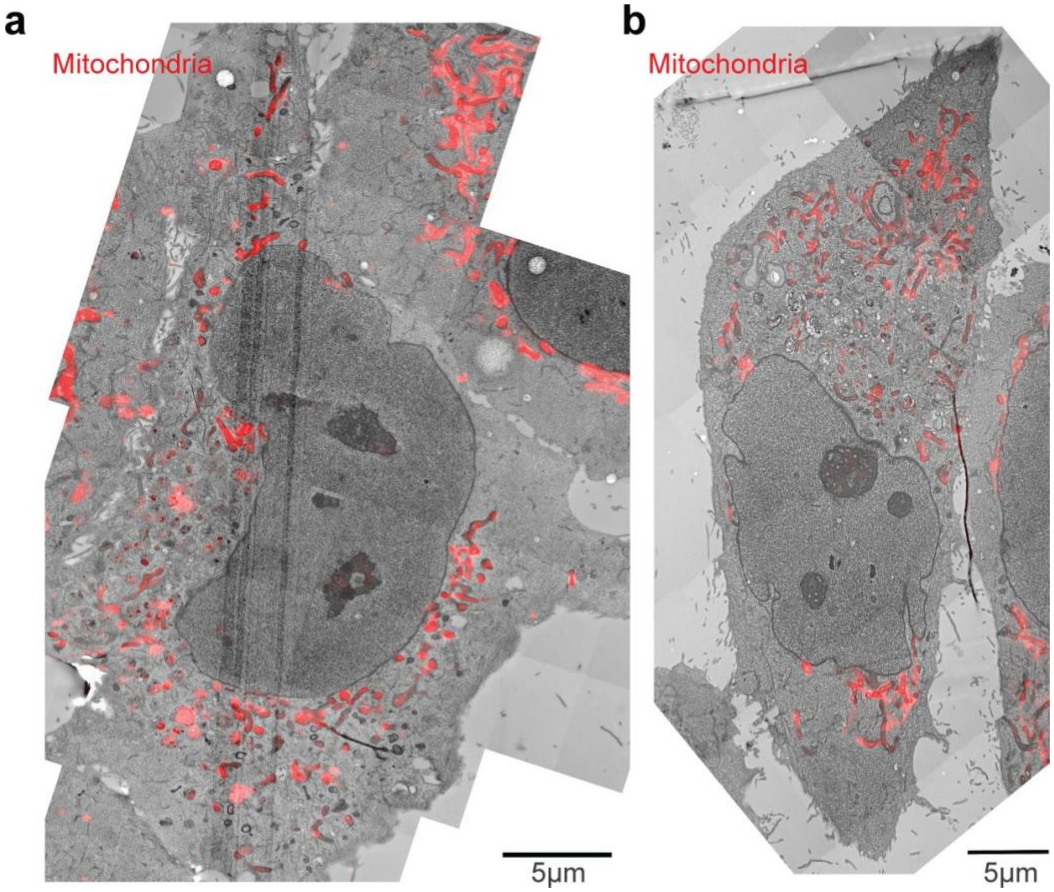


**Fig. S9. Correlation and re-registration in the z-dimension. a**, Dextran-labelled HeLa cell from Fig. 3d-f with the overlay of the MitoTracker Orange signal showing accurate fitting in z between TEM and 3D confocal microscopy. **b**, EGF-labelled HeLa cell from Fig. 3g-i with the overlay of the MitoTracker Orange signal showing accurate fitting in the z-dimension between TEM and 3D confocal microscopy.

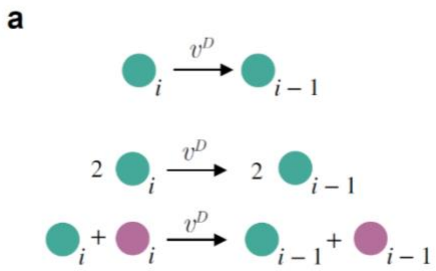


**Fig. S10. Combinations of hops possible with monomers and dimers of dyneins (green) and kinesins (magenta)**. We assumed that the hopping rate for pairs of motors is the same as that of a single motor. For an antagonistic pair made from a dynein and a kinesin motor, the hops can occur either to the right or to the left with rates $vD/\Delta x$ and $vK/\Delta x$ respectively.

### Supplementary Movie Legends

**Movie S1. Dynein binding events.** Cell expressing mDHC-GFP imaged using HILO microscopy. The green arrowhead points to a dynein molecule that became visible (a binding event), started moving towards the minus end of MT and disappeared (an unbinding event). Similarly, the magenta arrowhead points to a dynein molecule that bound and moved to the MT plus ends and the white arrowhead points to a stationary dynein molecule. 'N' marks the position/direction of the nucleus. Scale bar: 5$\mu m$.

**Movie S2. Dynamics of p150.** Cell expressing mCherry-p150 imaged using SD confocal microscopy. Unlike dynein whose association with the MT is short lived, p150 punctae are visible for long durations. The green arrowhead points to a p150 puncta moving towards the minus end of MT, presumably as part of the tripartite complex. The magenta arrowhead points to a p150 puncta moving away from the nucleus likely at a MT plus end. A stationary p150 spot indicated by the white arrowhead is visible for the entire duration of the movie. Note that the p150 signal is faint because a cell expressing low level of mCherry-p150 was chosen to avoid overexpression artefacts of p150. 'N' marks the position/direction of the nucleus. Scale bar: 5$\mu m$.

**Movie S3. Dextran is in Rab5+ compartments.** Cell expressing mCherry-Rab5 (green) and after taking up A647 tagged 10kDa dextran (magenta) imaged using SD microscopy. The white arrowheads point to dextran and Rab5 that colocalised for the duration of the video, indicating that dextran could be used as a marker for early endosomes. Scale bar: 5$\mu m$.

**Movie S4. Cargo-MT interaction.** Cell expressing mCherry-tubulin (green), post uptake of A647-tagged 10kDa dextran (magenta) imaged using SD microscopy. The white arrowheads point to dextran vesicles which remained close to the MTs even when stationary. Scale bar: 5$\mu m$.

**Movie S5. Intracellular crowding prevents cargo diffusion.** Cell post uptake of A647-tagged 10kDa dextran treated with 10$\mu M$ nocodazole to depolymerize the MTs. As expected, directed movement of the dextran vesicles was completely abrogated. The diffusion of the vesicles was also insignificant indicating that intracellular crowding likely prevented cargo diffusion away from the MTs. 'N' marks the position/direction of the nucleus. Scale bar: 5$\mu m$.

**Movie S6. Movement of dextran vesicles in HeLa cells.** Cell post uptake of A647-tagged 10kDa dextran imaged using HILO microscopy. Scale bar: 5$\mu m$.

**Movie S7. Movement of EGF vesicles in HeLa cells.** Cell post uptake of A647-tagged EGF imaged using HILO microscopy. Scale bar: 5$\mu m$.

**Movie S8. Activation of dynein upon dextran vesicle binding.** Cell expressing mDHC-GFP (green), post uptake of A647-tagged 10kDa dextran (magenta) imaged using HILO microscopy. The white arrowhead points to a dextran vesicle onto which a dynein molecule bound (indicated by an abrupt appearance of intensity in the GFP channel) followed by movement of the dynein-cargo complex towards the minus end of MTs. 'N' marks the position/direction of the nucleus. Scale bar: 5$\mu m$.

**Movie S9. Activation of dynein upon EGF vesicle binding.** Cell expressing mDHC-GFP (green), post uptake of A647-tagged EGF (magenta) imaged using HILO microscopy. The white arrowhead points to an EGF vesicle onto which a dynein molecule bound (indicated by an abrupt appearance of intensity in the GFP channel) followed by movement of the dynein-cargo complex towards the minus end of MTs. 'N' marks the position/direction of the nucleus. Scale bar: 5$\mu m$.

**Movie S10. EGF endosomes undertake large movements towards the nucleus.** Cell post uptake of A647-tagged EGF (magenta) imaged using HILO microscopy over a $\sim$17 min time period. 'N' marks the position/direction of the nucleus. Scale bar: 5$\mu m$.
